# CLIPB4 is a central node in the protease network that regulates humoral immunity in *Anopheles gambiae* mosquitoes

**DOI:** 10.1101/2023.07.07.545904

**Authors:** Xiufeng Zhang, Shasha Zhang, Junyao Kuang, Kathleen A. Sellens, Bianca Morejon, Sally A. Saab, Miao Li, Eve C. Metto, Chunju An, Christopher T. Culbertson, Mike A. Osta, Caterina Scoglio, Kristin Michel

## Abstract

Insect humoral immune responses are regulated in part by protease cascades, whose components circulate as zymogens in the hemolymph. In mosquitoes, these cascades consist of <u>c</u>lip domain <u>s</u>erine <u>p</u>roteases (cSPs) and/or their non-catalytic <u>h</u>omologs (cSPHs), which form a complex network, whose molecular make-up is not fully understood. Using a systems biology approach, based on a co-expression network of gene family members that function in melanization and co-immunoprecipitation using the serine protease inhibitor (SRPN)2, a key negative regulator of the melanization response in mosquitoes, we identify the cSP CLIPB4 from the African malaria mosquito *Anopheles gambiae* as a central node in this protease network. *CLIPB4* is tightly co-expressed with *SRPN2* and forms protein complexes with SRPN2 in the hemolymph of immune-challenged female mosquitoes. Genetic and biochemical approaches validate our network analysis and show that CLIPB4 is required for melanization and antibacterial immunity, acting as a prophenoloxidase (proPO)-activating protease, which is inhibited by SRPN2. In addition, we provide novel insight into the structural organization of the cSP network in *An. gambiae*, by demonstrating that CLIPB4 is able to activate proCLIPB8, a cSP upstream of the proPO-activating protease CLIPB9. These data provide the first evidence that, in mosquitoes, cSPs provide branching points in immune protease networks and deliver positive reinforcement in proPO activation cascades.

## Introduction

The African malaria mosquito, *Anopheles gambiae*, is one of the efficient vectors of life-threatening human malaria parasites. In 2021, approximately 247 million cases and 619,000 deaths were reported of malaria worldwide (1). Despite the availability of an effective, albeit non-sterilizing, vaccine, vector population control remains the most common and effective malaria prevention tool, which is threatened by insecticide resistance in target mosquito populations. Novel vector population control schemes, utilizing gene drive, male sterility, and *Wolbachia*-based methods (2–4) in part rely on detailed molecular understanding of mosquito physiology. For this reason, mosquito innate immunity has received much attention, as it is a major determinant of vector competence and survival of mosquitoes within and across their life stages (5–7).

Protease cascades play a central role in innate immunity, regulating blood clotting and complement activation in mammals, hemolymph clotting in horseshoe crabs, and melanization and Toll signaling pathway activation in insects (8–10). Their activation leads to proteolytic cleavage of a series of inactive protease zymogens between a prodomain and the protease domain, which then cleave downstream key immune factors, ensuring a fast, often within minutes, activation of antimicrobial immunity. In insects, these cascades consist of so-called clip-containing serine proteases (cSPs), a large protein family (consisting of 5 subfamilies, A-E), which is characterized by one or more clip pro-domains and one or more protease domains with trypsin-like specificity, and their homologs (cSPHs), which have lost one or more residues of the catalytic triad required for proteolytic activity. Our knowledge of the molecular makeup of these cascades is mainly derived from insect model species that are either large in size, enabling biochemical investigations of their hemolymph (e.g. *Manduca sexta* (11) and *Tenebrio molitor* (12)), or hold an excellent genetic and molecular tool box (in case of *Drosophila melanogaster* (13,14)).

At their core, canonical insect immune protease cascades consist of a modular serine protease (ModSP), which upon recognition of foreign molecular patterns, self-activates and then cleaves and activates a cSP in the subfamily C that in turn cleaves and activates a terminal cSP in the subfamily B (cSP-B) (15–17). Activation of melanization, an insect specific immune response proceeds through the terminal cSP-B pro-phenoloxidase activating protease (PAP) that cleaves the zymogen pro-phenoloxidase (proPO) to active phenoloxidase, a key enzyme in the conversion of aromatic amino acids to quinones that ultimately polymerize to eumelanin on foreign surfaces. Clip serine protease homologs support the proPO cascade, as they are required for effective activation of PO on foreign surfaces (18–20). To prevent uncontrolled spread and over-activation of responses, serine protease inhibitors in the serpin family tightly control mammalian and arthropod immune protease cascades.

In *An. gambiae*, melanization is a readily selectable parasite resistance mechanism that garnered much attention in hopes of exploiting the molecular underpinnings of this resistance mechanism for malaria control purposes (21,22). In addition, the gene families encoding cSPs and cSPHs in *An. gambiae*, known as CLIPs, and proPOs have undergone large expansions, raising the question whether regulation of melanization is more complex than in other insects (23–26). To date, the melanization response in *An. gambiae* has been analyzed genetically using RNAi (27) combined with at least five experimental readouts, often with the implicit or explicit assumption that identified CLIPs contribute biochemically to proPO activation. Thus far, two *An. gambiae* cSPs, CLIPB9 and B10 have been identified as PAPs, based on their ability to revert increased systemic melanization induced genetically by knockdown (kd) of *An. gambiae serpin-2* (*SRPN2*) and to cleave purified *M. sexta* proPO *in vitro* (28,29). These data indicate that, based on the canonical model of proPO activation cascades, melanization in *An. gambiae* depends on at least two such cascades, characterized by CLIPB9 and B10, respectively. Based on genetic and biochemical data, CLIPB8 is located further upstream of CLIPB9 (30). All three CLIPB proteases required for proteolytic activation of proPO are inhibited by serpins (SRPNs). Of the 18 SRPNs annotated in the *An. gambiae* genome, only SRPN2 has been shown to control systemic melanization (7,31). It inhibits effectively both PAPs, CLIPB9 and B10, and also CLIPB8, albeit to a lesser degree (28–30). Several additional CLIPBs, including CLIPB3, 4, 14 and 17 also contribute to melanization; however the molecular function and placement of these additional CLIPBs in proPO activation cascades is unknown (32,33). The only cSP in the CLIPC family to contribute to melanization is CLIPC9, presumably through the action of cleavage of either PAP, which however remains to be demonstrated (34).

In addition to cSPs, melanization in *An. gambiae* is regulated positively by a core module of cSPHs, consisting, in hierarchical order, of SPCLIP1, CLIPA8 and CLIPA28. This module is located genetically upstream of CLIPC9 and downstream of the thioester-containing protein TEP1 (6,32,34–36). Melanization is also regulated negatively by cSPHs, including CLIPA2 and A14 (32,37). While the precise mechanism(s) by which *An. gambiae* cSPHs regulate melanization (positively or negatively) are unknown, observations in *An. gambiae* suggest that cSPHs contribute to the opsonization of microbial surfaces by TEP1 and the leucine-rich repeat proteins LRIM1 and APLC1 (35,38), and, based on data from other insect species, to PO activity by mediating proPO cleavage by PAPs (19,39,40). In addition, melanization of rodent and human malaria parasites in *An. gambiae* is inhibited by two <u>c</u>-type lectins, CTL4 and CTLMA2 (41,42), which presumably bind to sugar moieties on the parasite surface and prevent melanization locally. The rodent malaria parasite melanization phenotype induced by *CTL4* kd has been used successfully to identify some of the cSPs and cSPHs mentioned above (e.g. (32)).

However, despite the significant number of studies summarized above, our understanding of cSP cascades that control melanization in *An. gambiae* remains incomplete, and more than 80% of the 110 annotated CLIPs await experimental exploration. To guide the experimental analysis of *An. gambiae* cSPs and to identify putative key regulators of the melanization response, we describe here a novel systems biology approach to study melanization in insects, combining transcriptomic network and proteomic analyses. Using this approach, we identify the cSP CLIPB4 as a central node in the protease network that regulates melanization in *An. gambiae* and characterize its placement in the proPO activation cascade using a combination of genetic and biochemical approaches.

## Materials and Methods

### Mosquito cultures

The *An. gambiae* G3 strain was obtained from the Malaria Research and Reference Reagent Resource Center (MR4) at CDC and reared as described previously (29). For egg production, females were provided a blood meal of commercial heparinized horse blood (Plasvacc USA) using parafilm as a membrane stretched across glass feeders connected to a 37°C circulating water bath.

### Anopheles gambiae Melanization Gene Co-expression Network (AgMelGCN)

The AgMelGCN was constructed according to (43). Genes of interest (GOIs), represented as nodes were selected based on published evidence either confirming experimentally their role in melanization or confirming phylogenetically their membership in protein families known to contribute to melanization in mosquitoes. The GOI list contained 191 genes, including 9 prophenoloxidases (PPO), 14 thioester-containing proteins (TEP), 23 C-type lectins (CTL), 16 leucine-rich repeat immune proteins (LRIM), 18 serpins (SRPN), 4 putative modular serine proteases (ModSP), 20 clip domain serine protease A family proteins (CLIPA), 29 CLIPB, 12 CLIPC, 13 CLIPD, 32 CLIPE, and SP24D, a putative ModSP. Their raw expression values were collated from 257 unique physiological conditions assessed in 30 separate transcriptome studies, and z-score conversion was performed for normalization under each condition. Two genes were considered co-expressed and connected by edges if the Pearson Correlation coefficient (PCC) of their expression vectors was above the 3% fitted sliding threshold. PCCs were rescaled based on the reversed curve of the fitted sliding thresholds to assign edge weights. The network was visualized using Gephi 0.9.2 (44) using the Fruchterman-Reingold force-directed algorithm (45). Network communities were assigned with the Girvan-Newman algorithm (46). Four measures were applied for node centrality, evaluating the edge abundance connected to a single node: node strength (sum of connected edge weights), eigenvector centrality, betweenness centrality and harmonic closeness centrality (47). In addition, *s*-core decomposition analysis was performed. In the case of a weighted network, the *s*-core subgraph consists of all nodes *i* with node strengths *s*(*i*) > *s*, where *s* is a threshold value. We define the threshold value of the s_n_-core as *s*_(n-1)_ = min *s*(*i*), among all nodes *i* belonging to the *s*_(n-1)-_core network. The *s_n-1_*-core is found by the iterative removal of all nodes with strengths *s*(*i*) ≤ *s*_(n-1)_ (48).

### SRPN2 Co-immunoprecipitation (Co-IP) and Mass Spectrometric Analysis

Female mosquitoes were challenged by microinjection of 69 nl lyophilized *Micrococcus luteus* in water (OD_600_ =0.55, Sigma-Adrich) per mosquito, and hemolymph was collected from 420 female mosquitoes into 1x Roche Protease Inhibitor in PBS by proboscis clipping 19 h post infection. The SRPN2 antiserum, produced previously (29), was affinity purified using the pH 7.2 coupling buffer with AminoLink^TM^ Plus Immobilization kit (ThermoFisher Scientific), and stored in PBS. Co-IP was performed with Pierce™ Protein A Magnetic Beads (ThermoFisher Scientific), and beads were rinsed with the buffer PBSOg (PBS with 0.7% w/v n-Octyl-BD-Glucopyranoside). Mosquito hemolymph was equally split for two co-IP reactions: one with 40 µl 1:1 slurry of PBSOg-containing magnetic beads cross-linked with 30 µg of purified SRPN2 antibody; and the other with 40 µl 1:1 slurry of PBSOg-containing control magnetic beads. Hemolymph was incubated with the beads for 1.5 h at room temperature with constant rotation. The unbound fractions were removed and magnetic beads were washed twice with PBSOg followed by four washes with UltraPure^TM^ water (Invitrogen). Bound proteins were eluted into 20 µl 0.1% TFA (w/v, trifluoroacetic acid in water) in each co-IP. Samples were shipped to the Nevada Proteomics Center (University of Nevada, Reno) for identification via LC-Orbitrap-MS. Contaminants were removed with 2-D Clean-up kit (GE Healthcare), and in-tube trypsin digestion was performed. Resultant peptides were run through a Michrom Paradigm Microdialysis Liquid Chromatography and Michrom CaptiveSpray coupled to a Thermo LTQ Orbitrap XL with Electron-transfer dissociation (ETD). MS spectra were analyzed using Sequest (IseNode in Proteome Discoverer version 2.2.0.388, Thermo Fisher Scientific). Sequest searches of ANOGA_uniprot_20210804.fasta (25565 entries) were performed with the following settings: digestion enzyme trypsin, maximal missed cleavage = 2, fragment ion mass tolerance = 0.60 Da, and a parent ion tolerance = 50 PPM. Carbamidomethyl of cysteine was specified as a fixed modification, while oxidation of methionine, acetylation of the N-terminus and phosphorylation of serine were specified as variable modifications. Scaffold (version 5.1.0, Proteome Software Inc.) was used to validate peptide and protein identifications. Peptide probability was calculated by the Peptide Prophet algorithm (49) with Scaffold delta-mass correction, and a ≥95.0% identification cut-off. Protein probabilities were assigned by the Protein Prophet algorithm (50). Proteins that contained similar peptides and could not be differentiated based on MS analysis alone were grouped to satisfy the principles of parsimony. In the SRPN2 Co-IP sample, a protein was considered present if protein probability was greater than 99.0%, and 2 minimal unique peptides were identified in the sequence. A protein was removed from the SRPN2 Co-IP list if at least one minimal peptide was detected in the control sample. Proteome Discoverer SequestHT scores were listed at the protein level to rank the final identification list.

### Expression and purification of recombinant proteins proCLIPB4 and proCLIPB4_Xa_

To produce recombinant proCLIPB4, its full coding region, including the signal peptide, was amplified by PCR using gene specific primers (Table S1) and *An. gambiae* adult cDNA as template. The forward primer contained a NotI site at 5’-end, and the reverse primer included three glycine codons, six histidine codons, a stop codon, and a HindIII site at the 3’-end. The PCR product was digested by NotI and HindIII, and inserted into the same restriction sites in the transfer vector pFastBac1 (Invitrogen). The resulting proCLIPB4-6His-pFastBac1 plasmid was used as a template to produce the mutant proCLIPB4_Xa_-6His-pFastBac1.To utilize commercial Factor Xa (New England Biolabs), the predicted activation site LTDR^107^ of proCLIPB4 was replaced with IEGR^107^. Site-directed mutagenesis was introduced to proCLIPB4-6His-pFastBac1 by Vent DNA polymerase (New England Biolabs) with mutagenic primers (Table S1), DpnI (New England Biolabs) digestion and NEB10β transformation. The resulting proCLIPB4_Xa_-6His-pFastBac1 was confirmed by Sanger Sequencing. The DNA segments containing proCLIPB4-6His and proCLIPB4_Xa_-6His, respectively, were transferred from pFastBac1 to pOET3 (Oxford Expression Technologies) by BamHI and HindIII sites, creating proCLIPB4-6His-pOET3 and proCLIPB4_Xa_-6His-pOET3, respectively. The pFastBac1 and pOET3 transfer vectors were used to produce corresponding CLIPB4 baculoviruses using the Sf9 cell line by means of Bac-to-Bac system (Invitrogen) or flashBAC-Baculovirus Expression system (Genway Biotech). Large-scale expression was performed using Sf9 suspension cultures at 2×10^6^ cells/mL cell density in Sf-900 II serum-free medium (Invitrogen). Baculoviruses were inoculated at multiplicity of infection (MOI) of 1, and expression media were harvested 48 h post infection. Recombinant proteins were purified as described previously (28–30). In brief, cells were cleared by centrifugation, and expression medium was dialyzed and subjected to column chromatography with Ni-NTA agarose (Qiagen) and Q Sepharose Fast Flow (Cytiva). Eluted fractions were screened by immunoblot with THE™ His Tag Antibody. Combined fractions were concentrated with buffer exchange (20 mM Tris, 50 mM NaCl, pH 8.0) in Amicon™ Ultra-10K centrifugal units. Recombinant proteins were stored at −80°C until future use. Recombinant proCLIPB8_Xa_, proCLIPB8, proCLIPB9_Xa_, proCLIPB9 and SRPN2 were expressed and purified as described previously (29,30).

### SDS-PAGE and Immunoblot

Protein samples were mixed with SDS sample buffer containing β-mercaptoethanol, and denatured at 95°C for 5 min. Proteins were separated on SDS-PAGE gels (Bio-Rad) with constant voltage at 120V followed by 180V. SDS-PAGE gels were stained with EZ-Run Protein Gel Staining Solution (Fisher) and de-stained with Milli-Q water. For immunoblot, proteins were transferred onto nitrocellulose membranes (GE Healthcare) at 10V constant voltage using Owl^TM^ Semidry Electroblotting Systems (ThermoFisher Scientific). Membranes were incubated with 5% dry milk in 1xTBST (0.05% Tween-20) at room temperature (RT) for 1 h, followed by primary antibody incubation diluted in 0.5% dry milk in 1xTBST (4°C overnight or RT for 2 h). Blots were washed three times with 1xTBST for 20 min each at RT. Membranes were further incubated with diluted secondary antibodies in 0.5% dry milk 1xTBST at RT for 1 h. Protein signals were developed by Alkaline Phosphate (AP) Conjugate Substrate Kit (Bio-Rad) and Amersham^TM^ ECL^TM^ Western Blotting Detection Reagents (GE Healthcare) for AP-conjugated and HRP-conjugated secondary antibodies, respectively. Images were captured by CCD camera of Azure Biosystems 300 for ECL development.

### Factor Xa activation of IEGR-mutated recombinant proteins

Factor Xa activation of recombinant proCLIPB4_Xa_ is described in the legend of online suppl. Fig. S3. Activation of proCLIPB8_Xa_ and proCLIPB9_Xa_ was performed as described previously (29,30).

### SRPN2 inhibition of CLIPB4

To produce active CLIPB4, 630 ng of recombinant proCLIPB4_Xa_ was incubated with 1 µg of Factor Xa in a total volume of 11 µl with Factor Xa activation buffer at 37℃ overnight. Two negative controls were set up in parallel, where proCLIPB4_Xa_ or Factor Xa was replaced with the same volume of Factor Xa activation buffer. Activated CLIPB4_Xa_ (154 ng/per reaction) was mixed with recombinant (r)SRPN2 at molar ratios of 1:1 or 1:10 (proCLIPB4_Xa_:SRPN2). After incubation at room temperature for 15 min, the reaction mixtures were subjected to immunoblot analysis with mouse anti-His, rabbit anti-SRPN2 or rabbit anti-CLIPB4 as primary antibodies. For Mass spectrometry identification, reactions were set up as described as above, and subjected to SDS-PAGE. After Coomassie brilliant blue staining, the band of interest (∼72 kDa) was excised. In-gel trypsin digestion and Electrospray ionization mass spectrometry (ESI-MS, Bruker Daltonics HCT Ultra) were performed at the Biotechnology/Proteomics Core Facility (Kansas State University). MS results were analyzed in Scaffold 4.0. To determine the stoichiometry of inhibition, 90 ng of Factor Xa-activated rCLIPB4_Xa_ was mixed with rSRPN2 at molar ratios of 0:1, 0.5:1, 1:1, 2:1, 4:1, and 6:1 (rSRPN2:rCLIPB4_Xa_) in reaction buffer 20 mM Tris, 150 mM NaCl, pH 8.0 supplemented with 3 µg BSA. Reactions were incubated at room temperature for 15 min. The residual amidase activity was determined with IEARpNA substrate at OD_405_ as described above. IEARase activity of Factor Xa was measured in parallel and subtracted from the above reactions of rSRPN2:rCLIPB4_Xa_. Amidase activity at molar ratio of 0:1 (rSRPN2:rCLIPB4_Xa_) was defined as 100%. All assays were performed in triplicate.

### Purification of PPO1&2 from M. sexta larval hemolymph

Prophenoloxidase 1&2 (*Ms*PPO1&2) was purified from *M. sexta* hemolymph similarly to published methodology (51). Briefly, hemolymph was collected into 100% ammonium sulfate (AS) at 0 °C from 5^th^ instar Day 2 *M. sexta* larvae. Fraction of 38-48% AS precipitation was dissolved in buffer A (10 mM KPi, pH 6.8, 500 mM NaCl, 0.5 mM reduced L-Glutathione), and dialyzed against buffer A. Bio-Gel HT Hydroxyapatite column (Bio-Rad) was utilized to separate proteins via a linear gradient of 10-100 mM KPi in buffer A. Two microliters per fraction was assayed with 200 µl 2 mM Dopamine hydrochloride in 50 mM NaPi, pH 6.5 with and without 0.1% cetylpyridinium chloride (CPC, w/v) in 96-well plates, and absorbance change was monitored at 470 nm in EPOCH 2 microplate reader (BioTek). Fractions containing CPC-activated PO activity were combined, and concentrations of MgCl_2_ and CaCl_2_ were adjusted to 1 mM each. The protein solution was applied twice through a Concanavalin A Sepharose 4B column (GE Healthcare), equilibrated in 20 mM Tris, pH 7.4, 0.5 M NaCl, 1 mM MgCl_2_, 1 mM CaCl_2_. Column flow-through fraction was adjusted to 1 M AS, and loaded onto a Phenyl Sepharose 6 Fast Flow low sub column (Cytiva), equilibrated in 0.1 M KPi, pH 7.1, 1 M AS. Elution was performed with a descending linear gradient of 0.1 M KPi, pH 7.1, 1 M AS to 0.01 M KPi, pH 7.1. Fractions with CPC-activated PO activity were pooled and concentrated to 1.1 ml using Amicon^TM^ Ultra-15 30K centrifugal units (Merck Millipore). Proteins were separated by Sephacryl S-300 column (Cytiva), equilibrated in 10 mM MOPS, pH 7.2, 0.5 M NaCl. Fractions containing CPC-activated PO activity were pooled and concentrated to 4.6 mg/ml by Amicon^TM^ Ultra-15 30K units. Equal volume of glycerol was mixed with purified *Ms*PPO1&2, and the purified zymogen was stored at −80°C at 2.3 mg/ml in 5 mM MOPS, pH 7.2, 0.25 M NaCl, 50% glycerol.

### ProPO activation by CLIPB4_Xa_ using M. sexta plasma and purified MsPPO1&2

To evaluate the contribution of CLIPB4 on proPO activity, *M. sexta* plasma and purified *Ms*PPO1&2 were utilized as described previously (29). Naïve hemolymph was collected from a cut proleg of 5^th^ instar Day 3 *M. sexta* larvae, and centrifuged at 5,000 g, 4℃ for 5 min to remove hemocytes and cell debris. Factor Xa was used to activate proCLIPB4_Xa_ as described in online suppl. Fig. S3. For PO enzymatic activity plate assay, in each well, 1 µl plasma was mixed with 3 µl Factor Xa-proCLIPB4_Xa_ activation mixture per well, supplemented to 10 µl with buffer B (20 mM Tris, 20 mM NaCl, 6.7 mM CaCl_2_, pH 7.5). Three negative controls were set up, replacing Factor Xa, proCLIPB4_Xa_, and both with buffer B, respectively for a final reaction volume of 10 µl. Three technical replicates were set up per reaction, and incubated at RT for 15 min. Per well, 200 µl 2 mM Dopamine hydrochloride was added in the buffer 50 mM NaPi, pH 6.5. Absorbance was monitored at 470 nm at RT for 1 h. For PO activity assay with purified MsPPO1&2, per well, 1 µg purified *Ms*PPO1&2 (4 µl at 250 ng/µl) was mixed with 5 µl of activation mixture and 3 µl of buffer B for a total reaction volume of 12 µl. One additional negative control was included by mixing 1 µg purified *Ms*PPO1&2 with 8 µl buffer B. Three technical replicates were set up for each reaction and incubated at RT for 1 h. Dopamine hydrochloride was added as described above and absorbance was monitored at 470 nm at RT for 1 h. One unit of PO activity was defined as ΔA470/min = 0.001.

### ProPO cleavage by CLIPB4_Xa_ using M. sexta plasma and purified MsPPO1&2

To detect cleavage of *Ms*PPO1&2, we performed immunoblots with rabbit anti-*Ms*PPO1&2 primary antibody (51). Naïve *M. sexta* plasma was first diluted to 1:20 in 20 mM Tris, 150 mM NaCl, pH 8.0. Reactions were set up by mixing 1 µl diluted plasma, 3 µl Factor Xa activation buffer or 3 µl each from Factor Xa proCLIPB4_Xa_ RT activation mixtures (including negative controls), and 6 µl buffer B for a 10 µl total reaction volume. Three additional samples were prepared by mixing 3 µl each from Factor Xa proCLIPB4_Xa_ RT activation mixtures and 7 µl buffer B. Reactions were incubated at RT for 20 min. For purified *Ms*PPO1&2, reactions were set up by mixing 2 µl purified *Ms*PPO1&2 (250 ng/µl, 500 ng) with 6 µl Factor Xa activation buffer or 6 µl each from Factor Xa proCLIPB4_Xa_ activation mixtures (see online suppl. Fig. 3). Reactions were incubated at RT for 20 min. For immunoblot, 1.2 µl of each reaction (containing 75 ng of *Ms*PPO1&2) was mixed with 8.8 µl water, and boiled at 95°C for 5 min with SDS sample buffer. Samples were separated on 7.5% SDS-PAGE gels, and probed with *Ms*PPO1&2 antibody. In addition, CLIPB9_Xa_-*Ms*PPO1&2 reactions were included as a positive control. Briefly, to obtain activated CLIPB9_Xa_, 2.5 µg of purified recombinant proCLIPB9_Xa_ was mixed with 1 µg Factor Xa, and Factor Xa activation buffer in a total reaction volume of 25 µl, and incubated at 37°C overnight. Two negative control reactions were set up and incubated in parallel, in which recombinant proCLIPB9_Xa_ and Factor Xa were replaced with the same volume of Factor Xa activation buffer, respectively.

### CLIPB4 activation of proCLIPB8 and proCLIPB9

To test whether CLIPB4 cleaves recombinant proCLIPB8 and/or proCLIPB9, proCLIPB4_Xa_ was activated by Factor Xa at 37℃. 154 ng of Factor Xa-activated CLIPB4_Xa_ was incubated with 100 ng proCLIPB8 or 100 ng proCLIPB9 at RT for 15 min. Recombinant proCLIPB8_Xa_ or proCLIPB9_Xa_ was activated by Factor Xa similarly as described previously (29,30), and 100 ng of activated CLIPB8_Xa_ or CLIPB9_Xa_ was used as a positive control. Samples were subjected to SDS-PAGE and immunoblot using mouse anti-His, rabbit anti-CLIPB8, or rabbit anti-CLIPB9 antibodies. To measure the enzymatic activity of cleaved CLIPB8, 610 ng of recombinant proCLIPB4_Xa_ (102 ng/µl) was incubated at 37 ℃ for 10 h with 1 µg (1 µg/µl) Factor Xa, supplemented to 21 µl with Factor Xa activation buffer. 72 ng of activated CLIPB4_Xa_ was incubated with 450 ng proCLIPB8 (225 ng/µl) at room temperature for 1 h. Reactions were supplemented with 200 µl of 100 µM N-benzoyl-Phe-Val-Arg-p-nitroanilide (FVR-pNA, EMD Chemicals) in 0.1 M Tris-HCl, pH 8.0, 0.1 M NaCl, 5 mM CaCl_2_ and amidase activity was measured continuously at OD_405_ for 30 min. 450 ng of activated CLIPB8_Xa_ was used as a positive control. One unit of CLIPB8 activity was defined as ΔA405/min = 0.001. All activity assays were performed in triplicate.

### Double-stranded RNA (dsRNA) synthesis and RNAi experiments

DsRNA synthesis and adult female mosquito injections were performed as described previously (29). Primers are listed in Table S1 for T7-tagged gene-specific amplification of *CLIPB4*. Primers were adapted from previous studies for dsRNA synthesis of *SRPN2*, *GFP*, *CLIPB8*, and *LacZ* (29,30,52). In single gene knockdown (kd) experiments, 3 to 4-day old adult female mosquitoes were injected each with 207 ng of dsRNA in a total volume of 69 nl. For kd control, mosquitoes were injected with the same quantity of the non-related dsRNA of either *Green Fluorescent Protein* (ds*GFP*) or *β-galactosidase* (ds*LacZ*). For double knockdowns (dkd), mosquitoes were injected with 138 nL of a 1:1 dsRNA mixture containing 1.5 µg/µl of each dsRNA. For dkd controls, ds*GFP* was added to compensate the total dsRNA dose to 414 ng/mosquito.

### Melanization phenotype analysis

To assess CLIPB4’s impact on melanization, two assays were utilized, Melanization-associated Spot Assay (MelASA) and SRPN2 depletion-induced melanotic pseudotumor formation. To quantify melanotic excreta, MelASA was performed in the standard format (50 mosquitoes/experimental cup) as described previously (34), with or without bacterial challenge. For MelASAs after bacterial challenge, dsRNA-injected mosquitoes were allowed to recover for 60 h and injected each with 50.6 nl of 1x PBS, OD_600_=1.5 *Escherichia coli* (resuspended in 1x PBS, same below), OD_600_=1.5 *Staphylococcus aureus*, and OD_600_= 0.4 *M. luteus* (*Micrococcus lysodeikticus* lyophilized cells ATCC, No. 4698, Sigma - Aldrich), respectively. Clean P8-grade white filter papers (Fisher) were inserted into the bottom of the experimental cups after bacterial challenge and removed 12 h later. For MelASAs without bacterial challenge, clean filter papers were inserted immediately after dsRNA injection, and removed 60 h later. All MelASAs were performed with five independent biological replicates using different mosquito generations. Collected filters papers were imaged under white epi-illumination without filters in an Alpha Imager System, and analyzed by Fiji is Just Image J (FIJI, Version 2.1.0) (53) using custom FIJI macros. To quantify melanotic pseudotumor formation, mosquitoes were dissected when mortality reached 70% in the ds*SRPN2*/ds*GFP* group. Abdominal walls were prepared as described previously (7) and examined under 40× magnification with Axio Imager A1 microscope (Zeiss) equipped with AxioCam MR5 (Zeiss). Image processing, and abdomen melanized area calculation were performed in ImageJ software as described previously (29,54). Five independent biological replicates were performed using 20 mosquitoes per treatment. All statistical analyses were performed using GraphPad Prism (ver. 9.5.1, GraphPad Software).

## Results

### Co-expression analysis of putative melanization genes identifies CLIPB4 as a central node

In the absence of a genome-wide protein-protein interaction network for *An. gambiae*, we used gene co-expression as a proxy to identify proteins that are likely to inhabit the same space at the same time. To facilitate the identification of possible protein-protein interactions, and specifically SRPN2-cSP interactions controlling melanization, we therefore constructed an undirected gene co-expression network of all *An. gambiae* genes belonging to gene families experimentally shown to contribute to melanization, which we named the *<u>A</u>nopheles <u>g</u>ambiae* <u>Mel</u>anization <u>G</u>ene <u>C</u>o-expression <u>N</u>etwork (AgMelGCN) (Fig. 1a). In total, the AgMelGCN contains 178 nodes (genes) and 1229 edges (co-expression links between gene pairs) (online suppl. Tables S2 and S3). To reveal the central nodes of AgMelGCN, we performed an *s*-core analysis (*s*=11.14). The core contained 25 genes, of which nine play a role in melanization (APL1C, CLIPA2, A8, B4, B10, CLIPC9, CTL4, CTLMA2, LRIM1; online suppl. Table S2). To identify key cSPs in the network, we calculated two centrality measures, node strength (weight sum of all node’s edges) and harmonic closeness centrality (sum of shortest paths to all other nodes within the full network) (47). Of all annotated cSPs, CLIPB4 was the highest ranked node in either centrality measure (online suppl. Table S2, Fig. 1b). In AgMelGCN, CLIPB4 was connected by a weighted edge to 44 other nodes, including seven SRPNs; notably, the SRPN2-CLIPB4 edge had the highest weight among all SRPN nodes connected to CLIPB4 (Fig. 1b; online suppl. Table S3). SRPN2 was connected to 18 nodes, including five cSPs, all in the CLIPB sub-family (Fig.1c; online suppl. Table S3), of which the SRPN2-CLIPB4 edge possessed the highest weight. Overall, these results suggested that CLIPB4 is a central node in the AgMelGCN whose expression most closely resembled that of SRPN2.

**Fig. 1.**
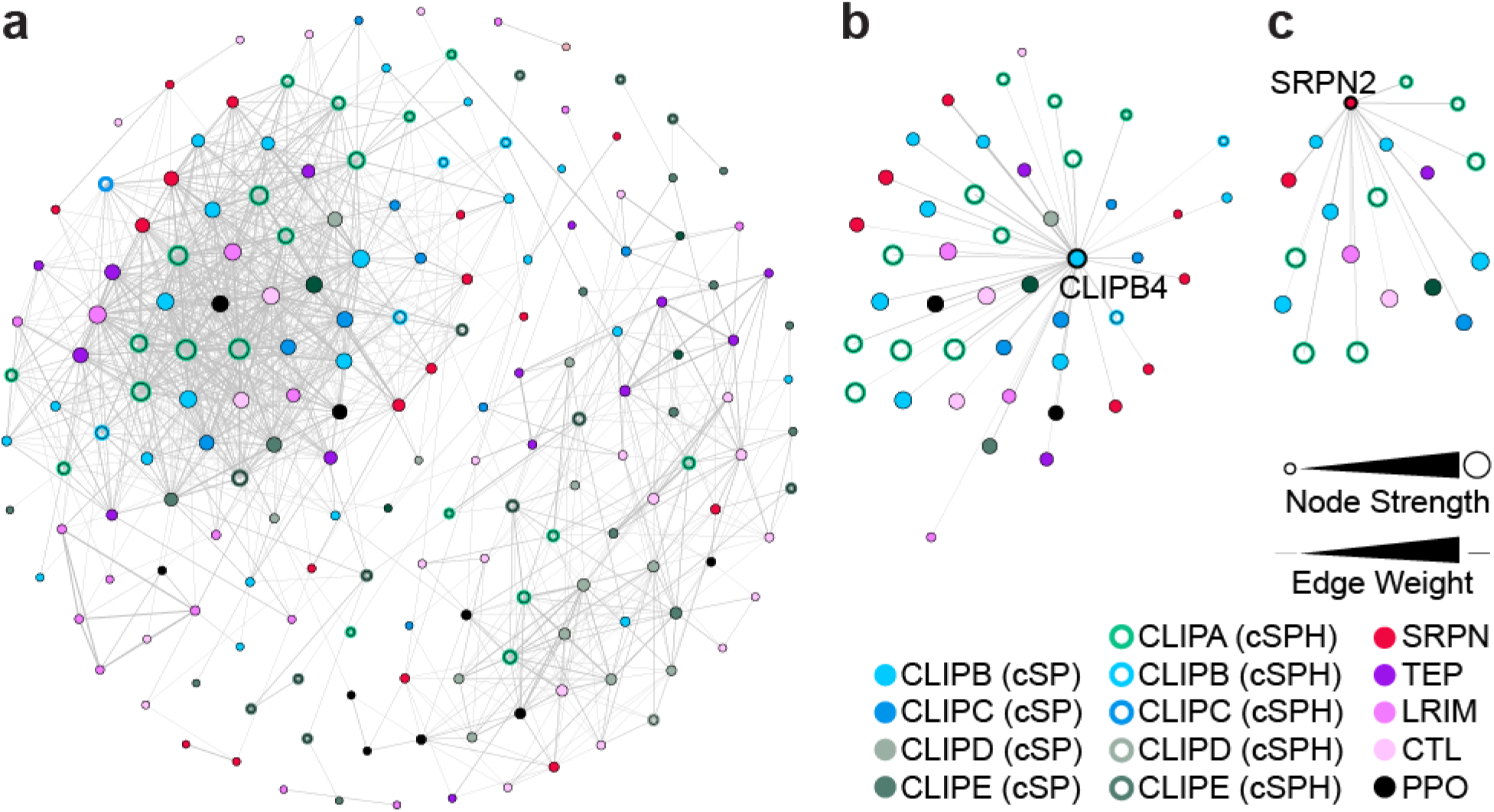
Gene co-expression network of genes with putative roles in melanization (AgMelGCN). (A) The network topography of AgMelGCN containing 178 nodes and 1229 edges. Of all annotated putative CLIP serine proteases, CLIPB4 has the highest node degree in the network and is co-expressed with SRPN2. (B) Genes in the network that are co-expressed with CLIPB4, which includes SRPN2. (C) Genes in the network that are co-expressed with SRPN2, which includes CLIPB4. The position of every node in the network visualization is identical in all three panels.

### SRPN2 and CLIPB4 interact in adult mosquito hemolymph

Since SRPN2 is a critical factor affecting the melanization process, we aimed to identify its closely associated proteins in the hemolymph of adult mosquitoes. Using co-immunoprecipitation, we identified 21 proteins bound directly or indirectly to SRPN2 in the hemolymph of adult female mosquitoes 24 h post challenge with lyophilized *M. luteus* bacteria (Table 1). Among these, six genes were co-expressed with SRPN2 in AgMelGCN, including TEP15, CLIPB4, CLIPA14, SRPN11, CLIPA9 and CLIPA6 (Fig. 1c, Table 1 and online suppl. Table S3). Among annotated cSPs and cSPHs, CLIPB4 was ranked the highest in SRPN2 Co-IP MS protein scores, warranting further examination of the interaction between SRPN2 and CLIPB4.

**Table 1.**
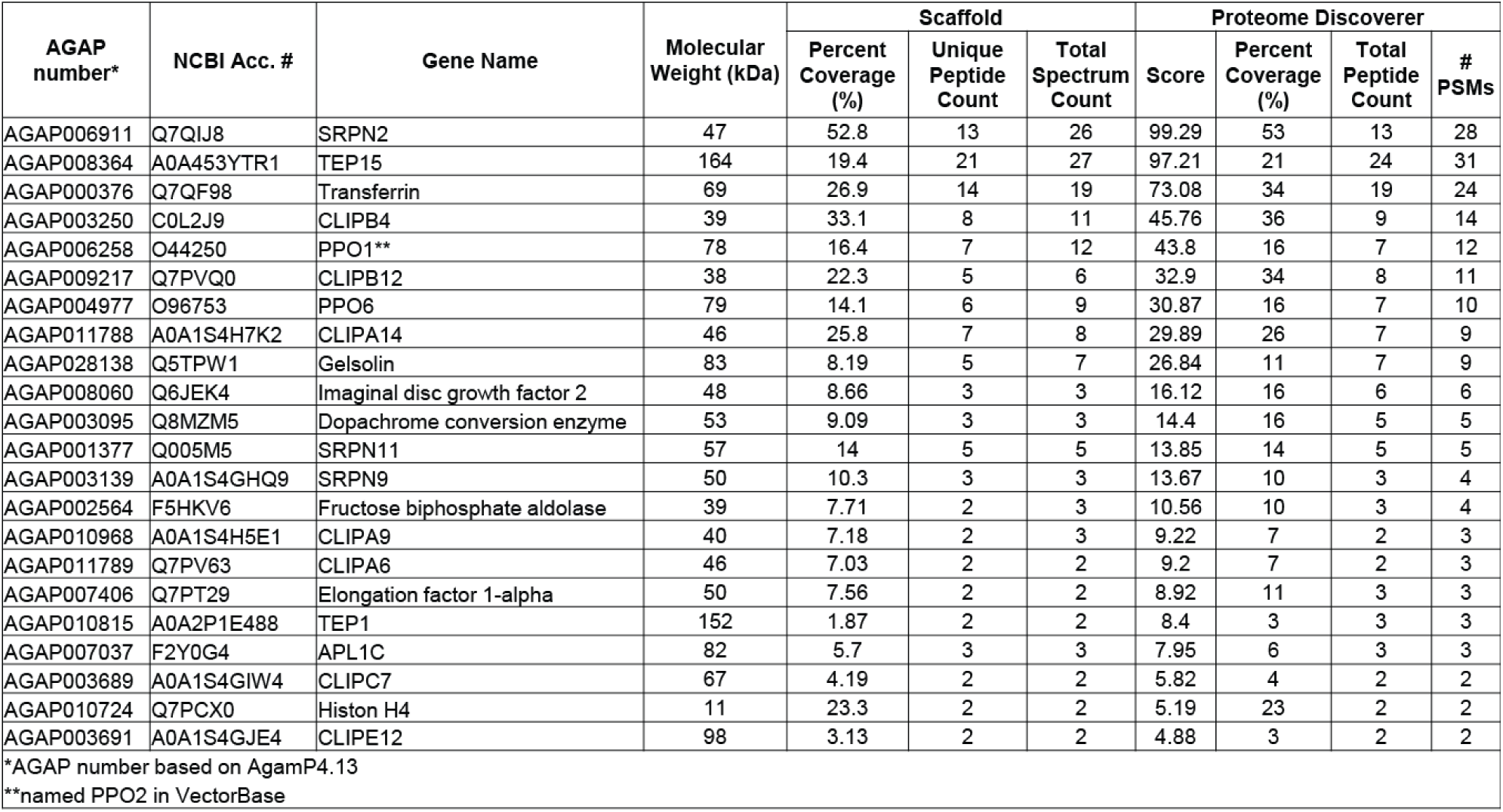
List of proteins that co-immunoprecipitate with SRPN2.

### CLIPB4 is a serine protease with trypsin-like specificity

To investigate its biochemical function(s), we expressed recombinant *An. gambiae* proCLIPB4 zymogen *in vitro*. The determinants of enzyme specificity were predicted to be D^305^, G^332^ and G^343^ (online suppl. Fig. S1a), suggesting that CLIPB4 is a trypsin-like protease that cleaves its substrate after arginine or lysine residues (55). As the endogenous activating enzyme of CLIPB4 is unknown, a mutant (proCLIPB4_Xa_) was produced by replacing its putative activation site ^104^LTDR^107^(56) with ^104^IEGR^107^ (online suppl. Fig. S1b), enabling recognition and cleavage by commercially available bovine Factor Xa. We produced and purified recombinant proCLIPB4_Xa_ zymogen (rproCLIPB4_Xa_) tagged with 6His at its C-terminus (Fig. S2a). Coomassie stain, His antibody and anti-CLIPB4 antibody detected rproCLIPB4_Xa_ (37.9 kDa) at the expected molecular weight. Recombinant proCLIPB4_Xa_ was cleaved by Factor Xa after incubation at 37℃ and RT, generating a fragment corresponding to the expected size of the CLIPB4 catalytic domain (rCLIPB4_Xa_, Fig. S3a and c). Recombinant CLIPB4_Xa_ exhibited amidase activity, using IEAR*p*NA substrate, while rproCLIPB4_Xa_ did not (Fig. S3b and S3d). As noted in previous works, Factor Xa also utilized IEAR*p*NA as a substrate (28–30). These results together support that, consistent with our construct design, Factor Xa is able to cleave and activate rCLIPB4_Xa_-6His.

### SRPN2 irreversibly inhibits CLIPB4 in vitro

Based on the AgMelGCN and co-IP results, we hypothesized that SRPN2 is a direct inhibitor of CLIPB4. To test this hypothesis, we determined (i) if SRPN2 and CLIPB4 form inhibitory protein complexes and (ii) if SRPN2 decreases CLIPB4 amidase activity in a concentration-dependent manner. Purified rSRPN2, tagged N-terminally with 6-His tag, was incubated with rCLIPB4_Xa_ at room temperature for 15 min at two different molar ratios, 1:1 or 10:1 (rSRPN2: rCLIPB4_Xa_). In SDS-PAGE and immunoblot analyses of these reactions a higher molecular weight band appeared at 72 kDa, which was detected with anti-His, anti-SRPN2 and anti-CLIPB4 antibodies (Fig. 2A, 2B and 2C). ESI-MS analysis on the 72 kDa band (online suppl. Table S4) detected 15 and 2 peptides for SRPN2 and CLIPB4, respectively, corresponding to 11 and 2 unique peptides for each, confirming that SRPN2 and CLIPB4 form a SDS-stable protein complex. CLIPB4_Xa_ amidase activity against IEAR*p*NA decreased with increasing amount of rSRPN2, with a stoichiometry of inhibition (SI) of 1.4 (Fig. 2d), demonstrating that SRPN2 functions as an efficient inhibitor of CLIPB4.

**Fig. 2.**
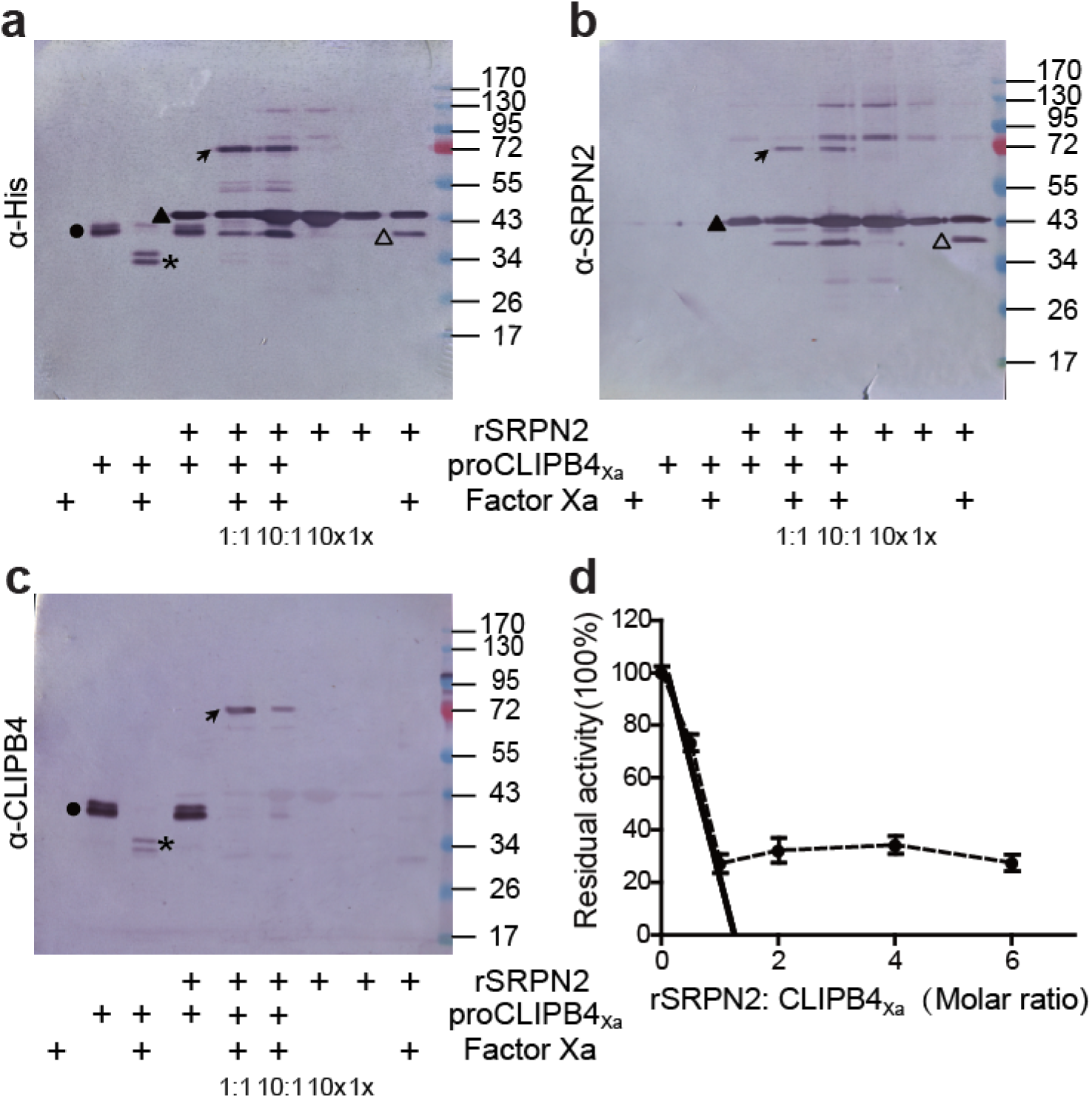
SRPN2 forms SDS-stable complex with CLIPB4 and inhibits its activity. (**a**, **b**, **c**) Detection of CLIPB4-SRPN2 complex by immunoblot analysis. Recombinant proCLIPB4_Xa_ was activated by Factor Xa, and then incubated with recombinant SRPN2 with a molar ratio of 1:1 or 10:1 (rSRPN2: proCLIPB4_Xa_) at room temperature for 15 min. The mixtures were subjected to 12% SDS-PAGE followed by immunoblot analysis with mouse anti-His (**a**) or rabbit anti-SRPN2 (**b**) or rabbit anti-CLIPB4 (**c**) antibodies. Circles, the proCLIPB4_Xa_ zymogen; asterisks, catalytic domain of proCLIPB4_Xa_; solid triangles, recombinant SRPN2; hollow triangles, truncated SRPN2; *arrows*, CLIPB4/SRPN2 complex. (D) Stoichiometry of SRPN2 inhibition of CLIPB4. Factor Xa-activated CLIPB4 was incubated with recombinant SRPN2 at various molar ratios at room temperature for 15 min. Residual activities of CLIPB4 were measured with IEAR*p*NA as substrate and plotted as mean±1 SD (n=3) against the corresponding molar ratios of rSRPN2 and proCLIPB4_Xa_.

### CLIPB4kd is required for the phenotypes induced by SRPN2 depletion

Previously, *SRPN2*kd was shown to reduce the longevity of female mosquitoes and to induce melanotic pseudotumor formation (7). Our biochemical data support that SRPN2 directly inhibits the activity of CLIPB4, and we hypothesized that CLIPB4 is required for the phenotypes induced by SRPN2 depletion. To test this hypothesis, we performed genetic epistasis experiments, inducing gene knockdown by long dsRNA injection. Adult female *An. gambiae* were injected with dsRNAs targeting *CLIPB4*, *SRPN2*, and both *CLIPB4* and *SRPN2*, respectively. Injection of *dsGFP* was used as a control. *CLIPB4* knockdown was significant and specific as determined by immunoblot (online suppl. Fig. S4). Injection of ds*SRPN2* did not affect CLIPB4 protein levels.

We first evaluated the effect of *CLIPB4*kd on survival comparing the survival curves between the four different treatment groups, and we also monitored the survival of uninjected mosquitoes. *SRPN2*kd significantly shortened longevity as compared to the uninjected controls and ds*GFP*-control treatment (Fig. 3a and b). In contrast, *CLIPB4*kd had no effect on lifespan as compared to the controls. Importantly, ds*CLIPB4/SRPN2*-treatment substantially and significantly increased mosquito longevity as compared to *SRPN2*kd mosquitoes, returning longevity to those of uninjected and ds*GFP*-control treated mosquitoes.

**Fig. 3.**
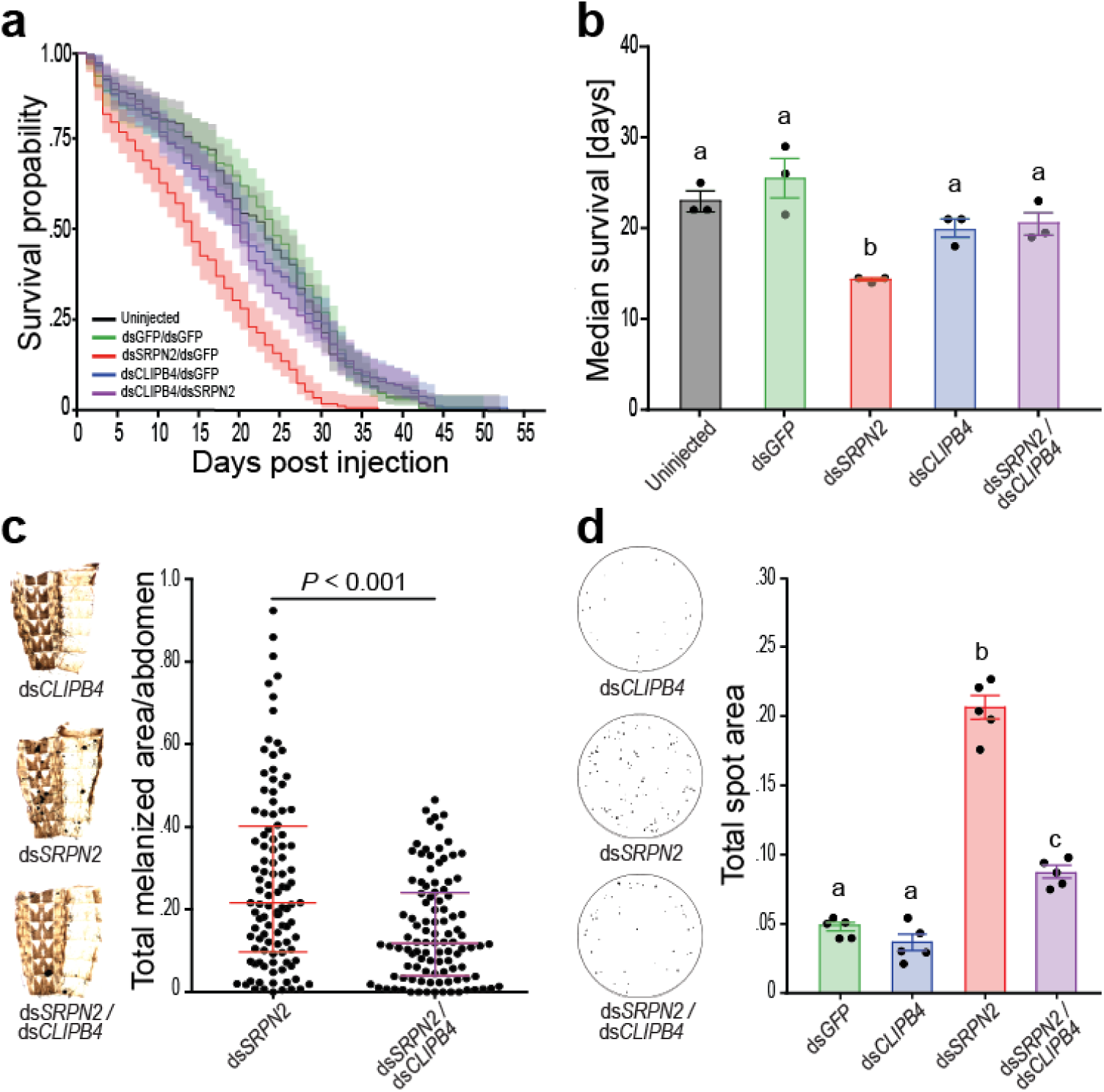
ds*CLIPB4* partially recovers the phenotype of SRPN2 depletion. (**a**) *CLIPB4*kd alone did not affect survival rates, but reverted the increased mortality induced by SRPN2 depletion. Survival curves were plotted as means with 95% confidence intervals (n=3). (**b**) Median survivals in days were plotted (mean ± 1 SEM) based on results shown in (**a**); statistically significantly differences are indicated by different letters (one-way ANOVA followed by Newman-Keuls test, P<0.01). (**c**) Total melanotic area per abdomen decreased due to ds*CLIPB4* in SRPN2-depleted genetic background. Medians and interquartile ranges are marked (n=104, 5 biological replicates), and statistical analysis was performed with the Mann-Whitney U test. (**d**) SRPN2 depletion increased melanotic excreta, which was tempered partially by ds*CLIPB4* treatment. Data are shown as dots for individual replicates, bars represent means ± 1 SD (n = 5); Data were distributed normally (Shapiro-Wilk test, *P*<0.0001), and statistically significantly differences are indicated by different letters (one-way ANOVA followed by Newman-Keuls test, P<0.01).

We then evaluated the effect of *CLIPB4*kd on melanization, using the appearance of melanotic tumors as a read-out (Fig. 3c). Injection of ds*GFP* or ds*CLIPB4* alone did not cause melanization (data not shown). In contrast, *CLIPB4*kd in a *SRPN2*-depleted background, significantly affected the *SRPN2*kd melanization phenotype, reducing the average total melanotic area for each dissected mosquito abdomen by 54 %, however, not completely reverting the SRPN2 depletion phenotype.

We also used the recently established melanization-associated spot assay (MelASA) to evaluate the contribution of CLIPB4 on the SRPN2 depletion phenotype (34). The standard format (50 mosquitoes/cup) was adopted and melanotic excreta were recorded for the four treatment groups. *CLIPB4*kd alone exhibited similar melanotic excreta area compared to that of the negative control, ds*GFP*. SRPN2 depletion, however, significantly increased melanotic excreta of female mosquitoes. Double knockdown of *SRPN2* and *CLIPB4* significantly reduced the excreta compared to *SRPN2* single knockdown, but did not completely abolish the SRPN2 depletion phenotype (Fig. 3d). Taken together, the silencing experiments strongly suggest that *CLIPB4* is epistatic to *SRPN2* and therefore downstream of this serpin in the proPO activation cascade.

### CLIPB4 is required for melanization after microbial challenge

Since CLIPB4 is required for melanization induced by SRPN2 depletion, we hypothesized that CLIPB4 is broadly required for melanization in antimicrobial immunity. To test the hypothesis, we first conducted MelASAs after challenge with bacteria (*S. aureus*, *M. luteus*, and *E. coli*) (online suppl. Fig. S5). Without bacterial challenge, only small amounts of melanotic excreta were observed in the ds*GFP-*injected and *CLIPB4*kd treatment groups, consistent with our previous observation in Fig. 3d. Each bacterial challenge increased significantly melanotic excreta in the ds*GFP*-injected group as compared to ds*GFP*-injected mosquitoes treated with PBS, supporting that bacterial treatment indeed induces the melanization immune response. The induction of melanotic excreta by bacterial challenge was dampened in the *CLIPB4*kd treatment groups. Independent of bacterial species, melanotic excreta were reduced by about 50% in *CLIPB4*kd mosquitoes as compared to ds*GFP*-injected mosquitoes, demonstrating that melanization in response to bacterial infection is broadly dependent on CLIPB4.

### CLIPB4 functions as a proPO activating enzyme

The shortage of hemolymph collectable from *An. gambiae* mosquitoes can be overcome by using the *M. sexta* model system as a source of proPO, as *M. sexta* PPO1&2 share the conserved cleavage site with eight of the nine proPOs encoded in the *An. gambiae* genome (7). Activated CLIPB4_Xa_ was incubated with *M. sexta* plasma, followed by immunoblot analysis using anti-*M. sexta* proPO antibody. A doublet band around 80 kDa in *M. sexta* plasma represents heterodimeric proPO consisting of 79-kDa proPO1 and 80-kDa proPO2 (51). Addition of activated CLIPB4_Xa_ to *M. sexta* plasma resulted in the appearance of a 70-kDa doublet band corresponding to *M. sexta* active PO (Fig. 4a). The same doublet band was observed in the plasma supplemented with activated CLIPB9_Xa_, which we identified previously as a functional PAP in *An. gambiae* (29). PO activity of plasma increased in the presence of active CLIPB4_Xa_ (Fig. 4c). These results confirm that the proteolytic activity of CLIPB4 promotes proPO cleavage and PO activity. To determine the placement of CLIPB4 in the proPO activation cascade in *An. gambiae*, we next tested whether CLIPB4 can directly cleave and activate proPO. ProPO1&2 was purified from 5^th^ instar *M. sexta* larvae (Fig. S2c). By Coomassie Blue staining and immunoblot analysis with MsPPO antibody, a dimer was observed around 80 kDa, consistent with previous observations of purified full length *M. sexta* proPO1 and 2 (51). We then incubated Factor Xa-activated rCLIPB4_Xa_ with purified *M. sexta* proPO, using again rCLIPB9_Xa_ as a positive control. Neither rproCLIPB9_Xa_ zymogen nor Factor Xa alone were able to cleave MsPPO1&2, while active rCLIPB9_Xa_ cleaved purified MsPPO1&2 to produce the PO1&2 band at 70 kDa (Fig. 4b). Similarly, neither rproCLIPB4_Xa_ zymogen nor Factor Xa alone cleaved PPO1&2, but active rCLIPB4_Xa_ cleaved proPO1&2 (Fig. 4b), leading to an increase in PO activity (Fig. 4d), demonstrating that CLIPB4 functions as a PAP.

**Fig. 4.**
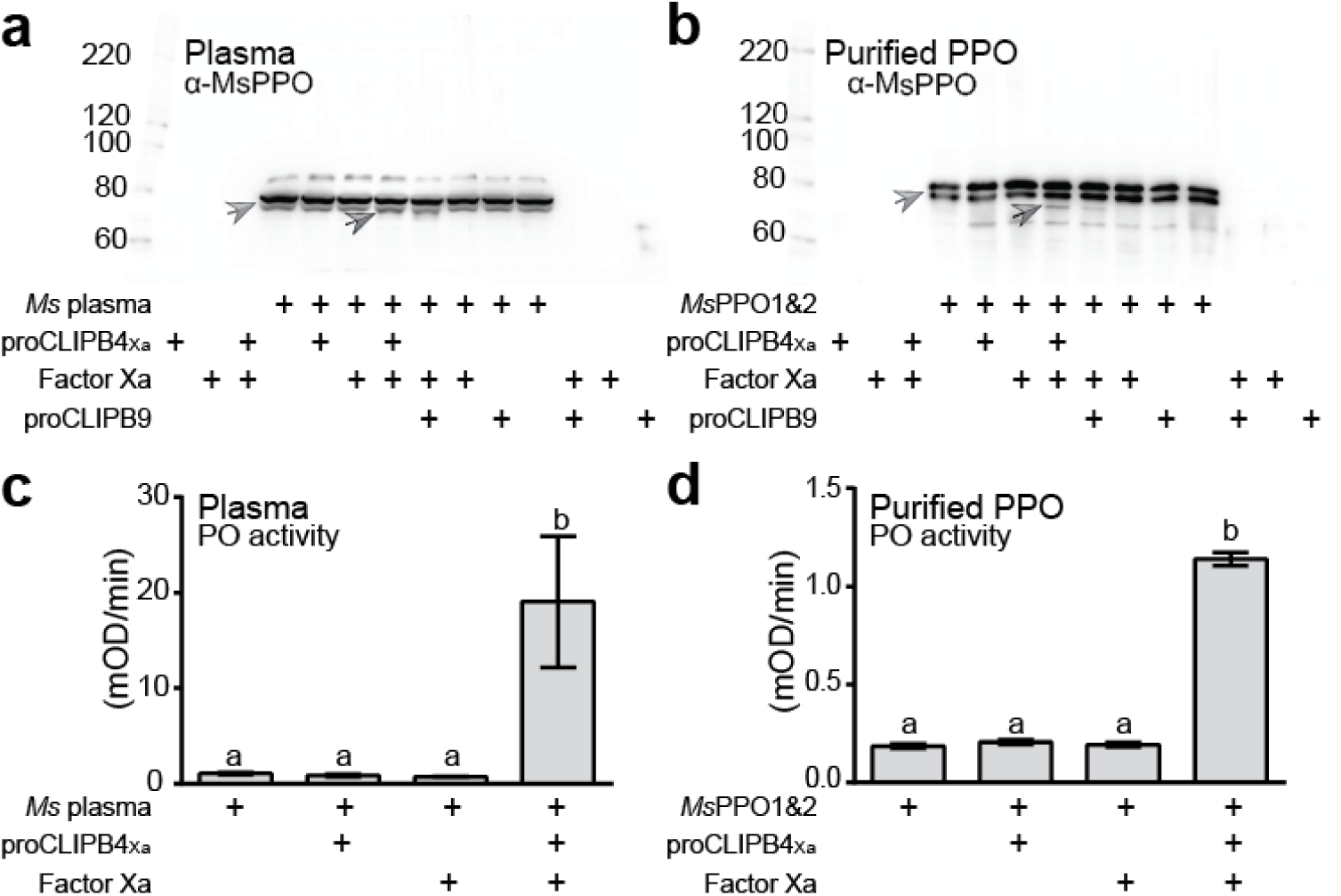
CLIPB4 cleaves and activates *M. sexta* PPO1&2. Factor Xa-activated proCLIPB4_Xa_ cleaves MsPPO1&2 in plasma (**a**) and purified protein (**b**), detected by immunoblot using rabbit anti-MsPPO1&2 antibody. The light and dark gray arrows indicate full and cleaved MsPPO1&2, respectively. CLIPB9 cleavage of *M. sexta* PPO was used as a positive control. The cleavage by CLIPB4 leads to increased PO activity in plasma (**c**) and with purified PPO protein (**d**). Bars represent mean ± 1 SD (n = 3); statistically significantly differences are indicated by different letters (one-way ANOVA followed by Newman-Keuls test, P<0.05).

### CLIPB4 cleaves and activates CLIPB8

We next explored whether CLIPB4 also promotes PO activity indirectly by activating the PAP zymogen proCLIPB9 (29) and/or proCLIPB8, a protease upstream of CLIPB9 in the melanization cascade in *An. gambiae* (30). To test whether CLIPB4 can cleave proCLIPB8 and/or B9 *in vitro*, active CLIPB4_Xa_ was incubated with wild-type recombinant proCLIPB8 and proCLIPB9, respectively, followed by immunoblot analysis using either anti-His antibody or anti-CLIPB8 and anti-CLIPB9 antibodies, respectively. In the reaction of active CLIPB4_Xa_ with proCLIPB8 zymogen, anti-His and anti-CLIPB8 antibodies detected a 36 kDa band, corresponding to the CLIPB8 protease domain, which was also detected in the control reaction of proCLIPB8_Xa_ incubated with Factor Xa (Fig. 5a and b). In contrast, addition of active CLIPB4_Xa_ (or Factor Xa) did not result in the cleavage of proCLIPB9 (Fig. 5c and d), suggesting that CLIPB4 is not directly upstream of CLIPB9. To evaluate whether cleavage of recombinant proCLIPB8 by CLIPB4_Xa_ results in CLIPB8 activation, we measured the amidase activity of the reaction mixtures using FVR as a substrate (30). Each of the five control reactions, containing proCLIPB8, proCLIPB8_Xa_, Factor Xa, proCLIPB4_Xa_, and proCLIPB4_Xa_/proCLIPB8, respectively did not exhibit amidase activity. Active CLIPB4_Xa_ showed limited amidase activity, which was increased threefold when active CLIPB4_Xa_ was incubated with proCLIPB8 prior to the addition of substrate (Fig. 5e).

**Fig. 5.**
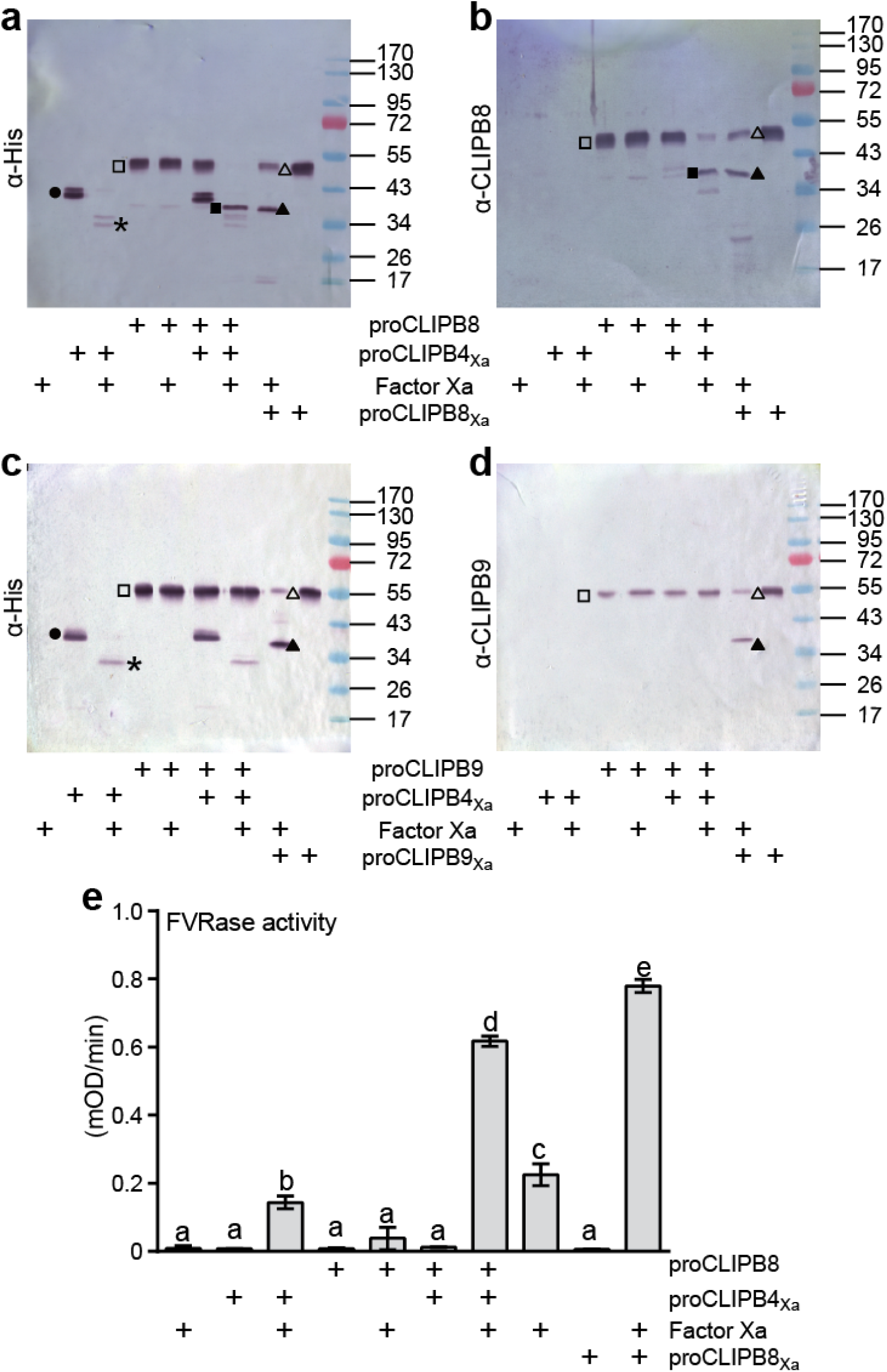
CLIPB4 activates B8, not B9. Recombinant proCLIPB4_Xa_ was activated by Factor Xa, and subsequently incubated with 100 ng of recombinant proCLIPB8 or proCLIPB9 at room temperature for 15 min. Samples were subjected to 12% SDS-PAGE followed by immunoblot with mouse anti-His (**a**, **c**), rabbit anti-CLIPB8 (**b**), and rabbit anti-CLIPB9 (**d**) antibodies, respectively. In parallel, 100 ng of Factor Xa-activated proCLIPB8_Xa_ or proCLIPB9_Xa_ were used as positive controls. Circles, proCLIPB4_Xa_ zymogen; asterisks, catalytic domain of CLIPB4; hollow squares, recombinant proCLIPB8 or proCLIPB9 zymogen; solid squares, catalytic domain of proCLIPB8; hollow triangles, proCLIPB8_Xa_ or proCLIPB9_Xa_ zymogen; solid triangles, catalytic domain of proCLIPB8_Xa_ or proCLIPB9_Xa_. (**e**) 72 ng of Factor Xa-activated proCLIPB4_Xa_ was incubated with 450 ng of recombinant proCLIPB8 at room temperature for 1 h, and the catalytic activity of CLIPB8 was determined using FVR as substrate; 450 ng of Factor Xa-activated proCLIPB8_Xa_ was used as a positive control. ars represent mean ± 1 SD (n = 3); statistically significantly differences are indicated by different letters (one-way ANOVA followed by Newman-Keuls test, P<0.05).

Together with the immunoblot results (Fig. 5a and 5b), FVRase activity assay supports that CLIPB4 specifically cleaves and activates CLIPB8.

To examine this phenomenon *in vivo*, ds*LacZ*, ds*CLIPB4* and ds*CLIPB8* were injected in adult female mosquitoes that were challenged by *S. aureus* post the dsRNA injection recovery. Hemolymph was collected 10 h after challenge and subjected to SDS-PAGE and immunoblot detection (Fig. 6). The Apolipophorin-II (APOII) was utilized as a loading control. CLIPB8 was depleted with ds*CLIPB8* treatment, serving as a positive control for CLIPB8 antibody detection. Ten hours after bacterial challenge, a clear reduction of CLIPB8 was observed in ds*LacZ* compared to the naïve control in two independent experiments, suggesting that CLIPB8 in the hemolymph was rapidly utilized in the acute immune response (57). The CLIPB8 reduction, mediated by *S. aureus* challenge, was rescued with ds*CLIPB4* treatment in two independent experiments, providing additional support that CLIPB4 acts upstream of CLIPB8 in the hierarchical cSP cascade regulating melanization in *An. gambiae*.

**Fig. 6.**
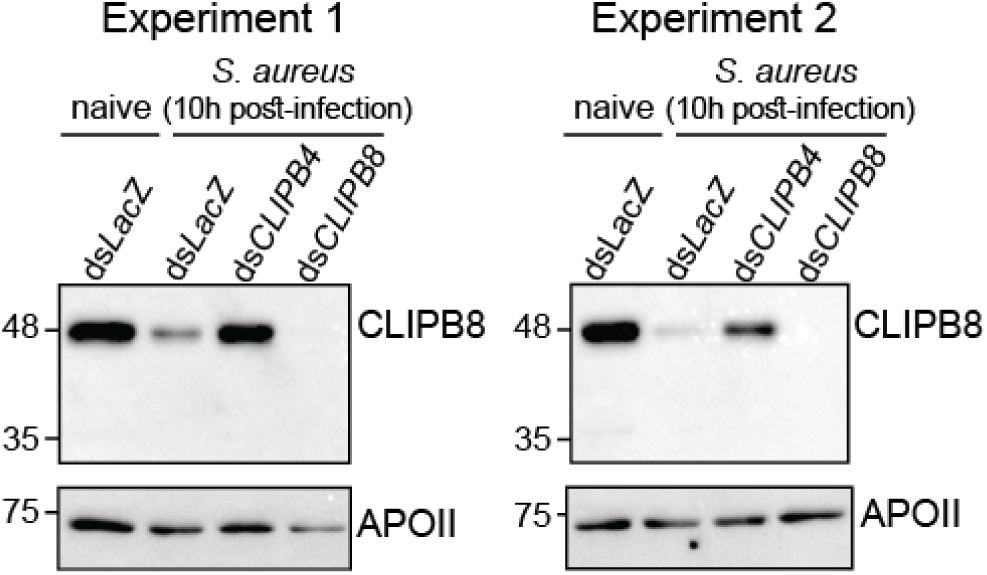
CLIPB4 is required for infection-induced depletion of CLIPB8 from the hemolymph of *An. gambiae*. Female mosquitoes were treated by ds*LacZ*, ds*CLIPB4* and ds*CLIPB8*. Posted recovery, mosquitoes were challenged by *S. aureus* (OD_600_=0.8). Ten hours post infection, hemolymph was collected by proboscis clipping, separated by SDS-PAGE, and subjected to immunoblot using rabbit anti-CLIPB8 and rabbit anti-APOII antibodies. Results were shown from two independent experiments.

## Discussion

Mosquitoes interact with a broad range of organisms across the symbiosis continuum and deploy melanization as a broad-spectrum immune defense against many organisms they encounter (58). The melanization of microbial surfaces in *An. gambiae* requires opsonization through TEP1 and the activity of PO to produce eumelanin, which depends on the action of cascades made up of cSPs and cSPHs. This study identifies CLIPB4 as an important factor required for melanization in response to microbial infection. CLIPB4 was previously shown to contribute to melanization of glass beads and malaria parasites in *CTL4* kd mosquitoes (32,33). Here we show that CLIPB4 is also required for melanotic tumor formation exerted by the depletion of SRPN2 in adult female mosquitoes, demonstrating that its contribution to melanization is not limited to specific surfaces, and suggesting that CLIPB4 is fundamental to eumelanin production. Indeed, our data demonstrate that CLIPB4 functions as a PAP, mediating proPO activation *ex vivo* and *in vitro*. In addition to CLIPB9 and B10, CLIPB4 is the third PAP identified in *An. gambiae* (28,29). In all three cases, their ability to cleave and activate proPO *in vitro*, was demonstrated using purified proPO1 and 2 from the hemolymph of *M. sexta*. Given that *An. gambiae* PPO5 and 6, the most abundant proPOs in the hemolymph of adult female *An. gambiae*, share the same activation cleavage site with *M. sexta* proPO1&2, it is highly likely that all three CLIPBs are bona fide PAPs in *An. gambiae* (7,59).

The existence of three PAPs, each constituting a terminal protease in a cSP cascade, suggests that melanization in adult female *An. gambiae* is regulated by at least three proPO activation cleavage cascades. Similarly, *M. sexta* proPO activation is regulated by at least two proPO activation cascades, utilizing either PAP3 or PAP1/PAP2 as the terminal proteases to cleave proPO (60,61). Likewise, two distinct proPO activation cascades regulate melanization in the lepidopteran species, *Helicoverpa amigera* (15,62). In *An. gambiae*, possibly not all three PAPs contribute equally to activate proPO. Expression of CLIPB4 in adult female *An. gambiae*, as measured by RNA-Seq is much higher than that of CLIPB9, B10, and any other CLIPB whose kd causes a melanization phenotype (Buchon and Michel, unpublished), suggesting that CLIPB4 protein, based on its abundance, contributes more to proPO activation than other PAPs. This observation may also explain why neither CLIPB8, B9 nor B10 were identified in the SRPN2 co-IP analyses, despite previous data demonstrating that CLIPB9 and SRPN2 form inhibitory complexes in *An. gambiae* hemolymph (29). Whether, in addition to protein concentration, the activation of the three *An. gambiae* PAPs depends on the type of microbial challenge, as observed in *H. amigera* (62), is currently under investigation.

In addition to its PAP function, CLIPB4’s central role in melanization in female *An. gambiae* is further manifested by its ability to cleave and activate proCLIPB8. This function was demonstrated *in vitro* by biochemical studies and *in vivo*, as *CLIPB4* kd reverts the consumption of CLIPB8 in the hemolymph of adult female *An. gambiae* upon microbial infection. This is of not only the first observation of multiple substrates for a single *An. gambiae* CLIP protease, but also the first demonstration of a CLIPB protease activating another proCLIPB zymogen in mosquitoes, breaking the mold of the classical hierarchy of CLIPC-CLIPB in insect immune protease cascades (63). The observation of CLIPB4 cleavage of CLIPB8 is paralleled in *M. sexta*, where PAP3 cleaves proHP5, a cSP-B family member (64). This cleavage of HP5, provides positive reinforcement of proPO activation, as HP5 in turn cleaves HP6, a cSP in the C family, upstream of PAP-1, linking the two proPO activation cascades in *M. sexta* (64). Based on these similar patterns in two evolutionarily distant insect species, it is tempting to speculate that positive reinforcement of proPO activation by cSP-B mediated activation of other cSP-Bs is a common theme in the melanization immune response.

Given that the regulation of melanization through proPO activation requires cSPs and cSPHs circulating in mosquito hemolymph prior to immune challenge, we posited that identification of important members in the cSP and cSPH cascades could be identified based on their co-expression. Indeed, the topology of the AgMelGCN, the co-expression network of members of gene families known to contribute to melanization clustered genes with known immune phenotypes into a single community. Using network science approaches, we then identified highly-connected nodes, referred to as hubs, which are integral to scale-free network stability (65). In gene co-expression networks, hub genes and are often relevant for network functionality, either as regulators or because they encode proteins with essential functions (66,67). The nodes with highest connectivity in the AgMelGCN encoded two cSPHs (CLIPA9 and 8), APLC1, and CLIPB4. All four genes encode proteins with known functions in innate immunity. *CLIPA9* kd in *An. gambiae* reduces *P. falciparum* oocysts, a phenotype that seems dependent on the presence of an unperturbed microbiota through an unknown mechanism (68). CLIPA8 is a key positive regulator of melanization and part of the cSPH cascade upstream of CLIPC9 (34,36). CLIPA8 is required for melanization of malaria parasites after *CTL4* kd, filamentous fungi, and bacteria (32,69,70). The leucine-rich repeat protein APLC1 contributes to antimicrobial immunity in mosquitoes by stabilizing TEP1 in the hemolymph, preventing its premature activation, and facilitating its delivery to microbial surfaces for opsonization (38,71,72). Likewise, CLIPB4 plays an important role in melanization through its integral roles in the cSP cascade that activates proPO described in this current study. Combined, these data support that gene co-expression networks can indeed be used to identify genes with important functions in mosquito innate immunity, and provide a blueprint for analyses of innate immunity in other insect species and likely other complex physiologies in mosquitoes.

The topology of the AgMelGCN also demonstrates the integration of the cSPH and cSP cascades through co-expression of members of each cascade. Our co-IP data indicate that these cascades also at least partially form protein complexes in the hemolymph of *An. gambiae*. We recently showed that CLIPB4 also cleaves CLIPA8 *in vitro*, which is required for CLIPA8 function (57), suggesting that the activation of cSPHs required for melanization is mediated by cSPs and potentially integrated with proPO activation through the formation of higher molecular weight protein complexes. Indeed, higher molecular weight complexes between cSPs and cSPHs and proPO have been described in other insect species, including *M. sexta* (73) and *Bombyx mori* (74), indicating that the formation of such immune complexes is likely an evolutionarily conserved process that is used to optimize proPO activity to microbial surfaces.

Based on our results presented herein and published data, we propose the following model of proPO activation in the melanization response of *An. gambiae* (Fig. S6). In adult female mosquitoes, circulating proPO zymogen is activated through the proteolytic activity of three CLIPB proteases, CLIPB4, B9 and B10 (28,29). CLIPB4 also activates proCLIPB8, which activates indirectly proCLIPB9 (30), providing positive reinforcement of proPO activation. Each of these CLIPBs provides an intervention point, as their activity is inhibited directly by SRPN2, reinforcing its role as the main negative regulator of melanization and survival in *An. gambiae* (7,31). This observation is paralleled in *M. sexta*, where serpin-3, the ortholog of SRPN2 inhibits all three identified PAPs (75,76). CLIPC9, currently the only known CLIPC protease required for melanization, is located likely upstream of one or more of the three PAPs and responsible for their proteolytic activation. In turn, based on the canonical proPO activation cascades in other insects, we hypothesize that proCLIPC9 is activated proteolytically through the action of an unknown ModSP. Additionally, melanization of microbial surfaces is further regulated through the activity of a cSPH cascade upstream of CLIPC9 (34,36). Whether, cSPHs also support melanization downstream of PAPs, as observed in other insect species (20,40,62) is currently under investigation. However, despite the existing complexity of this immune protease network, the presence of 110 CLIP genes in the *An. gambiae* genome suggests that additional CLIPB and CLIPC proteases may be involved (23,25,56). We are currently performing a reverse genetic screen to identify the contribution of all annotated *An. gambiae* cSPs and cSPHs to melanization and antimicrobial activity, leveraging the AgMelGCN and recently established *in vivo* bioassays and (34,77). Completion of this screen will describe the entire molecular make-up of the *An. gambiae* immune protease network and enable us to identify the molecular interactions within that are critical for vector competence and survival in this important vector species.

## Acknowledgements

We would like to thank all members of the Michel laboratory for mosquito rearing. We thank the former Michel laboratory members, Ms. Erin Peel for help with the reverse genetic experiments, and Dr. Xin Zhang for initial plasmid construction and expression of recombinant proteins. We are grateful to Dr. John Tomich and the members of the Biotechnology/Proteomics Core Facility, Kansas State University for the MS analysis of the SRPN2/CLIPB4 complexes. We thank Dr. Susan Paskewitz, University of Wisconsin for the kind gift of the CLIPB4 antibody. We gratefully acknowledge Ms. Rebekah J. Woolsey and Dr. David R. Quilici at the Mick Hitchcock, Ph.D. Nevada Proteomics Center Nevada Proteomics Center, University of Nevada, Reno, for providing the MS analysis of the co-IP experiments. Finally, sincere thanks got to Dr. Michael R. Kanost, Kansas State University, for his continued support and mentorship.

## Statement of Ethics

Not applicable to the enclosed manuscript.

## Author Contributions

K.M. conceived the study and C.A., C.T.C., M.A.O., C.S. and K.M. designed the experiments and supervised their execution. X.Z., S.Z., J.K., K.A.S., B.M. V., S.A.S., M.L., E.C.M., and K.M. performed the experiments and analyzed the data. X.Z., S.Z., and K.M. prepared the manuscript, and X.Z., S.Z., M.A.O., C.S., and K.M. revised the manuscript. All authors approved the submitted version.

## Data Availability

All generated or analyzed data are included within this article and its supplementary material files. Further enquiries should be directed to the corresponding author.

