## Supplementary Information for "CLIPB4 is a central node in the protease network that regulates humoral immunity in *Anopheles gambiae* mosquitoes"

**Supplementary Files**

**Online suppl. Table S1.** Oligonucleotides used in this study
**Online suppl. Table S2.** Node list, centrality measures, core and communities of AgMelGCN
**Online suppl. Table S3.** Undirected edge list and edge weight of AgMelGCN
**Online suppl. Table S4.** ESI-MS analysis of the CLIPB4-SRPN2 complex
**Online suppl. Fig. S1.** Sequence feature analysis of CLIPB4
**Online suppl. Fig. S2.** Recombinant and purified proteins in this study
**Online suppl. Fig. S3.** Activation of purified recombinant proCLIPB4$_{Xa}$ by Factor Xa
**Online suppl. Fig. S4.** ds*CLIPB4* injection depletes CLIPB4 in *An. gambiae* hemolymph
**Online suppl. Fig. S5.** *CLIPB4* is required for melanization after microbial challenge
**Online suppl. Fig. S6.** Model of ProPO activation in *An. gambiae*

**Online suppl. Table S1. Oligonucleotides used in this study**

*Amplification of full-length proCLIPB4, including signal peptide, with restriction sites for cloning in pFastBac1 vector:*

5’-ATGCGGCCGCATGATCGGTAATCGTGTGA-3’

5’-ACAAGCTTCTAATGGTGATGGTGATGATG*ACCTCCTCC*GTAGATATTGTCCTTTA-3’

Added *Not*I site is underlined.
Codon for three extra glysine is italicized.
Added *HindIII* site underlined.
The reverse complement sequence for 6 histidine residues inserted in front of the stop codon is double-underlined.

---

*CLIPB4 mutagenesis:*

CLIPB4 FT: 5’-TGCAGATCGAAGGACGTGTCATCGGCGGACAGCCGACCAAGA-3’
CLIPB4 FS: 5’-CGGCGGACAGCCGACCAAGA-3’
CLIPB4 RT: 5’-ATGACACGTCCTTCGATCTGCACCCCACAGTTTGGTGATTCT-3’
CLIPB4 RS: 5’-CCCCACAGTTTGGTGATTCT-3’
Sequence encoding IEGR instead of wild-type LTDR is underlined.

---

*CLIPB4 dsRNA synthesis:*

CLIPB4 dsRNA F: 5’-taatacgactcactatagggAGTGGGATCTTTCCTCGACC-3’
CLIPB4 dsRNA R: 5’-taatacgactcactatagggGCAAACAGATCGCGCTTAT-3’
Gene specific regions are in captial letters. T7 sequence is in lowercase letters.

### Online suppl. Table S2. Node list, centrality measures, core and communities of AgMelGCN

| AGAP# (Agam3.7) | Gene Name | Gene sub-family | Strength | Betweenness | Harmonic | Eigenvector | Core | Community |
|---|---|---|---|---|---|---|---|---|
| AGAP010968 | CLIPA9 | cSPH_A | 26.85093 | 0.034284 | 50.7655 | 0.214543 | core | 1 |
| AGAP010731 | CLIPA8 | cSPH_A | 26.47322 | 0.023305 | 50.2309 | 0.205371 | core | 1 |
| AGAP007033 | APL1C | LRIM | 24.18946 | 0.032678 | 47.9715 | 0.180722 | core | 1 |
| AGAP003250 | CLIPB4 | cSP_B | 23.90951 | 0.0599 | 48.8245 | 0.169625 | core | 1 |
| AGAP011791 | CLIPA1 | cSPH_A | 23.9073 | 0.010657 | 48.9617 | 0.199705 | core | 1 |
| AGAP000290 | CLIPA27 | cSPH_A | 23.69422 | 0.045262 | 48.7594 | 0.18084 | core | 1 |
| AGAP005334 | CTLMA2 | CTL | 22.96572 | 0.008346 | 48.385 | 0.194438 | core | 1 |
| AGAP003246 | CLIPB2 | cSP_B | 22.70925 | 0.014638 | 46.3453 | 0.180014 | core | 1 |
| AGAP006348 | LRIM1 | LRIM | 22.64712 | 0.004558 | 47.1986 | 0.189165 | core | 1 |
| AGAP011792 | CLIPA7 | cSPH_A | 22.63596 | 0.056882 | 48.1121 | 0.164304 | | 1 |
| AGAP009844 | CLIPB15 | cSP_B | 22.40939 | 0.037108 | 48.3707 | 0.190229 | core | 1 |
| AGAP005625 | SP213 | ModSP | 21.23292 | 0.016243 | 46.7821 | 0.171608 | core | 1 |
| AGAP004977 | PPO6 | PPO | 21.22101 | 0.006998 | 46.0017 | 0.182059 | core | 1 |
| AGAP004318 | CLIPC3 | cSP_C | 20.14973 | 0.02934 | 45.8955 | 0.157881 | core | 1 |
| AGAP029770 | CLIPB10 | cSP_B | 19.90973 | 0.001348 | 44.9828 | 0.16631 | core | 1 |
| AGAP005335 | CTL4 | CTL | 19.83408 | 0.001027 | 45.4705 | 0.178094 | core | 1 |
| AGAP004855 | CLIPB13 | cSP_B | 19.72421 | 0.00122 | 45.0983 | 0.168197 | core | 1 |
| AGAP011790 | CLIPA2 | cSPH_A | 19.22465 | 0.015729 | 45.3557 | 0.158836 | core | 1 |
| AGAP010816 | TEP3 | TEP | 18.53387 | 0.00321 | 44.3611 | 0.151625 | core | 1 |
| AGAP008654 | TEP12 | TEP | 18.45663 | 0.001412 | 43.5915 | 0.14342 | | 1 |
| AGAP011780 | CLIPA4 | cSPH_A | 18.33548 | 0.100539 | 45.0661 | 0.118357 | | 1 |
| AGAP004719 | CLIPC9 | cSP_C | 18.19304 | 0.054379 | 46.2382 | 0.159023 | core | 1 |
| AGAP028728 | CLIPE5 | cSP_E | 18.02949 | 0.000835 | 44.4492 | 0.156796 | core | 1 |
| AGAP012034 | CLIPC12 | cSP_C | 17.709 | 0.004366 | 44.8741 | 0.152125 | core | 1 |
| AGAP002813 | CLIPD6 | cSP_D | 17.61128 | 0.011877 | 43.9618 | 0.140197 | | 1 |
| AGAP012616 | PPO5 | PPO | 17.45024 | 0.011685 | 44.146 | 0.143712 | core | 1 |
| AGAP001377 | SRPN11 | SRPN | 16.84895 | 0.011492 | 44.4254 | 0.142953 | | 1 |
| AGAP006910 | SRPN3 | SRPN | 15.95843 | 0.012134 | 43.5304 | 0.134821 | core | 1 |
| AGAP028725 | SPCLIP1 | cSPH_A | 15.69849 | 0.001926 | 43.343 | 0.127089 | | 1 |
| AGAP010530 | CLIPE4 | cSPH_E | 15.42043 | 0.01971 | 42.868 | 0.133535 | core | 1 |
| AGAP007039 | LRIM4 | LRIM | 14.51691 | 0 | 40.9598 | 0.131898 | core | 1 |
| AGAP010545 | CLIPE10 | cSP_E | 13.97987 | 0.03454 | 42.6262 | 0.113632 | | 1 |
| AGAP010815 | TEP1 | TEP | 13.76325 | 0.019646 | 42.2238 | 0.107721 | | 1 |
| AGAP008364 | TEP15 | TEP | 13.62392 | 0.000514 | 42.2548 | 0.110917 | | 1 |
| AGAP003251 | CLIPB1 | cSP_B | 13.52545 | 0.003531 | 42.2841 | 0.107782 | | 1 |
| AGAP003057 | CLIPB8 | cSP_B | 12.91228 | 0.000899 | 41.5589 | 0.108063 | | 1 |
| AGAP008995 | CLIPD12 | cSP_D | 11.95417 | 0.020095 | 37.1973 | 0.000096 | | 0 |
| AGAP009221 | SRPN5 | SRPN | 11.80461 | 0.008731 | 40.4312 | 0.085721 | | 1 |
| AGAP006954 | CLIPA10 | cSPH_A | 11.60372 | 0.034348 | 37.2841 | 0.000109 | | 0 |
| AGAP009217 | CLIPB12 | cSPH_B | 11.24197 | 0.006741 | 40.0954 | 0.083253 | | 1 |
| AGAP003689 | CLIPC7 | cSPH_C | 11.02017 | 0.002953 | 39.7461 | 0.089198 | | 1 |
| AGAP012037 | CLIPB20 | cSP_B | 10.92501 | 0.003338 | 39.5406 | 0.093332 | | 1 |
| AGAP012502 | CLIPE25 | cSP_E | 10.5915 | 0.040575 | 36.2421 | 0.00009 | | 0 |
| AGAP008996 | CLIPD22 | cSP_D | 10.13188 | 0.011749 | 35.2003 | 0.000062 | | 0 |
| AGAP011788 | CLIPA14 | cSPH_A | 10.10792 | 0.002696 | 39.7546 | 0.072199 | | 1 |
| AGAP002270 | CLIPB7 | cSPH_B | 9.813112 | 0.000257 | 38.4063 | 0.080395 | | 1 |
| AGAP029559 | CTLMA6 | CTL | 9.651383 | 0.001798 | 34.8644 | 0.000108 | | 0 |
| AGAP006911 | SRPN2 | SRPN | 9.60437 | 0 | 39.0116 | 0.084333 | | 1 |
| AGAP010708 | CTLMA7 | CTL | 9.146309 | 0.019068 | 34.851 | 0.000032 | | 0 |
| AGAP001433 | CLIPD3 | cSP_D | 8.854533 | 0.013033 | 34.539 | 0.000041 | | 0 |
| AGAP001964 | CLIPA26 | cSPH_A | 8.748147 | 0.018041 | 36.1959 | 0.053809 | | 1 |
| AGAP009000 | CLIPD13 | cSP_D | 8.661522 | 0.003852 | 33.9995 | 0.00006 | | 0 |
| AGAP009316 | CTL10 | CTL | 8.654906 | 0.011171 | 35.2924 | 0.000074 | | 0 |
| AGAP011782 | CLIPE2 | cSPH_E | 8.581715 | 0.059194 | 37.1937 | 0.0009 | | 0 |
| AGAP008368 | TEP14 | TEP | 8.563013 | 0.007383 | 37.8411 | 0.054016 | | 1 |
| AGAP002815 | CLIPA15 | cSPH_A | 8.232504 | 0.006934 | 35.4847 | 0.000118 | | 0 |
| AGAP011793 | CLIPA31 | cSPH_A | 8.211923 | 0.013161 | 34.6394 | 0.000045 | | 0 |
| AGAP010830 | TEP9 | TEP | 8.209521 | 0.0868 | 37.0654 | 0.00042 | | 0 |

| AGAP# (Agam3.7) | Gene Name | Gene sub-family | Strength | Betweenness | Harmonic | Eigenvector | Core | Community |
|---|---|---|---|---|---|---|---|---|
| AGAP000573 | CLIPC4 | cSP_C | 8.109303 | 0.001926 | 38.81 | 0.052772 | | 1 |
| AGAP002825 | PPO1 | PPO | 7.999526 | 0.009309 | 33.955 | 0.00008 | | 0 |
| AGAP011787 | CLIPA5 | cSPH_A | 7.634444 | 0.000385 | 35.944 | 0.054685 | | 1 |
| AGAP007692 | SRPN14 | SRPN | 7.552971 | 0.007897 | 38.0236 | 0.050059 | | 1 |
| AGAP011789 | CLIPA6 | cSPH_A | 7.543558 | 0.001091 | 36.76 | 0.056653 | | 1 |
| AGAP012021 | CLIPE23 | cSPH_E | 7.500113 | 0.073575 | 37.4171 | 0.000962 | | 0 |
| AGAP001376 | SRPN17 | SRPN | 7.217464 | 0.006934 | 36.56 | 0.041408 | | 1 |
| AGAP010818 | TEP11 | TEP | 6.8518 | 0.009052 | 34.167 | 0.000033 | | 0 |
| AGAP010819 | TEP10 | TEP | 6.715981 | 0.010208 | 33.9504 | 0.000045 | | 0 |
| AGAP000315 | CLIPC6 | cSP_C | 6.518413 | 0.069723 | 38.5988 | 0.034869 | | 1 |
| AGAP009214 | CLIPB11 | cSP_B | 6.101442 | 0.008282 | 36.2449 | 0.017643 | | 1 |
| AGAP007037 | LRIM3 | LRIM | 6.09024 | 0 | 35.0415 | 0.039485 | | 1 |
| AGAP002784 | CLIPD8 | cSP_D | 6.070578 | 0.001027 | 31.5696 | 0.000018 | | 0 |
| AGAP004980 | PPO7 | PPO | 6.069014 | 0.049949 | 35.8356 | 0.000906 | | 0 |
| AGAP001375 | SRPN12 | SRPN | 5.949236 | 0.00122 | 29.1415 | 0.00002 | | 0 |
| AGAP004198 | SRPN19 | SRPN | 5.911029 | 0.01926 | 32.1382 | 0.00002 | | 0 |
| AGAP000940 | CTL7 | CTL | 5.883937 | 0.018939 | 32.7926 | 0.000038 | | 0 |
| AGAP008998 | CLIPD7 | cSP_D | 5.770825 | 0.001926 | 31.4298 | 0.000029 | | 0 |
| AGAP009211 | CLIPB42 | cSP_B | 5.667494 | 0.006613 | 31.0967 | 0.000035 | | 0 |
| AGAP011785 | CLIPE6 | cSPH_E | 5.629629 | 0.00841 | 36.1564 | 0.044436 | | 1 |
| AGAP001648 | CLIPB17 | cSP_B | 5.612322 | 0 | 35.6016 | 0.050673 | | 1 |
| AGAP009849 | CLIPB46 | cSP_B | 5.595291 | 0 | 34.7023 | 0.035522 | | 1 |
| AGAP008183 | CLIPD2 | cSP_D | 5.58847 | 0.005586 | 30.1942 | 0.000024 | | 0 |
| AGAP004975 | PPO3 | PPO | 5.581234 | 0.015152 | 30.6649 | 0.000073 | | 0 |
| AGAP011794 | CLIPA32 | cSPH_A | 5.526775 | 0.007383 | 32.861 | 0.000063 | | 0 |
| AGAP010812 | TEP4 | TEP | 5.400548 | 0 | 34.1196 | 0.03668 | | 1 |
| AGAP004149 | CLIPB41 | cSP_B | 5.396614 | 0.007062 | 31.8662 | 0.000071 | | 0 |
| AGAP011719 | CLIPE21 | cSP_E | 5.281665 | 0.003146 | 32.5469 | 0.000048 | | 0 |
| AGAP007454 | LRIM8A | LRIM | 5.234599 | 0.011941 | 33.3046 | 0.011924 | | 1 |
| AGAP028007 | CLIPC13 | cSP_C | 5.150178 | 0.069658 | 33.1968 | 0.000237 | | 0 |
| AGAP002911 | CTLMA9 | CTL | 4.835636 | 0.000642 | 29.2508 | 0.000012 | | 0 |
| AGAP002422 | CLIPD1 | cSP_D | 4.796412 | 0.012262 | 34.318 | 0.038923 | | 1 |
| AGAP003748 | CLIPE13 | cSP_E | 4.64503 | 0.003467 | 31.652 | 0.000049 | | 0 |
| AGAP002625 | CTL9 | CTL | 4.595335 | 0.010401 | 31.8449 | 0.000034 | | 0 |
| AGAP013089 | CLIPD20 | cSP_D | 4.564858 | 0.00886 | 32.2972 | 0.000073 | | 0 |
| AGAP007453 | LRIM9 | LRIM | 4.562108 | 0.009117 | 34.0579 | 0.01895 | | 1 |
| AGAP007693 | SRPN7 | SRPN | 4.363724 | 0.022535 | 30.3968 | 0.000164 | | 0 |
| AGAP013487-PA | CLIPB3a | cSP_B | 4.300922 | 0.089368 | 37.9411 | 0.0301 | | 1 |
| AGAP007457 | LRIM7 | LRIM | 4.288909 | 0 | 34.83 | 0.034102 | | 1 |
| AGAP009252 | CLIPE17 | cSPH_E | 4.156564 | 0.107601 | 36.9028 | 0.013391 | | 2 |
| AGAP011781 | CLIPA12 | cSPH_A | 4.106703 | 0 | 34.614 | 0.02147 | | 1 |
| AGAP007280 | SP212 | ModSP | 3.977559 | 0.000578 | 30.0968 | 0.000023 | | 0 |
| AGAP004976 | PPO8 | PPO | 3.950537 | 0.003082 | 29.393 | 0.000011 | | 0 |
| AGAP000123 | CTLSE2 | CTL | 3.937356 | 0 | 28.3285 | 0.000017 | | 0 |
| AGAP028167 | CLIPC14 | cSP_C | 3.912182 | 0.031266 | 31.9621 | 0.000377 | | 0 |
| AGAP010831 | TEP8 | TEP | 3.835757 | 0.007512 | 31.4041 | 0.000063 | | 0 |
| AGAP002811 | CLIPD4 | cSP_D | 3.829921 | 0.008539 | 29.9571 | 0.000044 | | 0 |
| AGAP003247 | CLIPB19 | cSP_B | 3.642661 | 0.011941 | 34.8715 | 0.021791 | | 1 |
| AGAP007456 | LRIM8B | LRIM | 3.575387 | 0.002119 | 32.4452 | 0.009951 | | 1 |
| AGAP009006 | CLIPD14 | cSPH_D | 3.458765 | 0.000128 | 28.1211 | 0.00001 | | 0 |
| AGAP003686 | CLIPB47 | cSP_B | 3.323784 | 0.024461 | 31.6029 | 0.003124 | | 1 |
| AGAP010832 | TEP19 | TEP | 3.314405 | 0.0165 | 30.7641 | 0.000114 | | 0 |
| AGAP007455 | LRIM10 | LRIM | 3.240873 | 0.000128 | 28.5669 | 0.002216 | | 1 |
| AGAP010628 | CLIPE30 | cSP_E | 3.116086 | 0.006484 | 31.2164 | 0.000047 | | 0 |
| AGAP010814 | TEP6 | TEP | 3.111733 | 0.003017 | 30.6679 | 0.000023 | | 0 |
| AGAP004148 | CLIPB5 | cSP_B | 3.098191 | 0 | 33.1057 | 0.014444 | | 1 |
| AGAP009213 | SRPN16 | SRPN | 3.032594 | 0.023819 | 34.7396 | 0.013752 | | 1 |
| AGAP013184 | CLIPB36 | cSPH_B | 2.995001 | 0.023498 | 33.8796 | 0.011397 | | 1 |
| AGAP006909 | SRPN1 | SRPN | 2.969735 | 0 | 32.0357 | 0.024826 | | 1 |
| AGAP006430 | CTLGA2 | CTL | 2.959753 | 0 | 28.1228 | 0.000014 | | 0 |

| AGAP# (Agam3.7) | Gene Name | Gene sub-family | Strength | Betweenness | Harmonic | Eigenvector | Core | Community |
|---|---|---|---|---|---|---|---|---|
| AGAP009670 | SRPN4 | SRPN | 2.88399 | 0.021957 | 31.2536 | 0.01788 | | 1 |
| AGAP007408 | CTLMA8 | CTL | 2.587849 | 0.023305 | 32.3766 | 0.000405 | | 0 |
| AGAP006631 | SP214 | ModSP | 2.583796 | 0.000449 | 28.1208 | 0.000038 | | 0 |
| AGAP011040 | CLIPE19 | cSP_E | 2.542688 | 0.000642 | 29.7033 | 0.000031 | | 0 |
| AGAP004978 | PPO9 | PPO | 2.53925 | 0.001156 | 30.2701 | 0.0096 | | 1 |
| AGAP007410 | CTLMA5 | CTL | 2.501595 | 0.004815 | 28.8497 | 0.000026 | | 0 |
| AGAP009273 | CLIPE18 | cSP_E | 2.4929 | 0.013482 | 31.4959 | 0.000369 | | 0 |
| AGAP009212 | SRPN6 | SRPN | 2.408935 | 0.024397 | 31.185 | 0.008554 | | 1 |
| AGAP010193 | CTLGA3 | CTL | 2.331348 | 0 | 30.4466 | 0.018077 | | 1 |
| AGAP009263 | CLIPB16 | cSPH_B | 2.299961 | 0 | 30.7287 | 0.016755 | | 1 |
| AGAP012020 | CLIPE22 | cSP_E | 2.171409 | 0.034027 | 27.9298 | 0.000009 | | 0 |
| AGAP028183 | CLIPE27 | cSPH_E | 2.065305 | 0.000706 | 25.7766 | 0.000003 | | 0 |
| AGAP007034 | LRIM11 | LRIM | 2.050424 | 0.000899 | 29.0506 | 0.000035 | | 0 |
| AGAP007045 | LRIM15 | LRIM | 2.028421 | 0.01849 | 33.7076 | 0.011123 | | 1 |
| AGAP000929 | CTLSE1 | CTL | 2.02494 | 0.011171 | 30.6385 | 0.000142 | | 0 |
| AGAP000572 | CLIPC10 | cSP_C | 2.017294 | 0 | 31.7537 | 0.012359 | | 1 |
| AGAP029769 | CLIPB9 | cSP_B | 1.979256 | 0.011043 | 30.8211 | 0.006643 | | 1 |
| AGAP003194 | SRPN8 | SRPN | 1.907742 | 0.023819 | 31.5853 | 0.004806 | | 1 |
| AGAP011325 | CLIPB45 | cSP_B | 1.906274 | 0.0052 | 27.8393 | 0.000014 | | 0 |
| AGAP029047 | CTL5 | CTL | 1.671559 | 0 | 27.3747 | 0.000036 | | 0 |
| AGAP011783 | CLIPA13 | cSPH_A | 1.602905 | 0.057974 | 33.199 | 0.005572 | | 1 |
| AGAP012938 | SRPN_x | SRPN | 1.598265 | 0.011043 | 23.148 | 0.000257 | | 1 |
| AGAP006258 | PPO2 | PPO | 1.595445 | 0.000257 | 25.5347 | 0.00004 | | 0 |
| AGAP012591 | CLIPA3 | cSPH_A | 1.583941 | 0 | 28.7563 | 0.000063 | | 0 |
| AGAP004810 | CTL3 | CTL | 1.51658 | 0 | 25.5569 | 0.000003 | | 0 |
| AGAP004811 | CTL1 | CTL | 1.510174 | 0.000128 | 25.6742 | 0.000004 | | 0 |
| AGAP008407 | TEP13 | TEP | 1.462239 | 0 | 25.8783 | 0.000007 | | 0 |
| AGAP012022 | CLIPE24 | cSP_E | 1.460037 | 0.000064 | 23.7198 | 0.000001 | | 0 |
| AGAP007407 | CTLMA4 | CTL | 1.456695 | 0 | 28.2936 | 0.005191 | | 1 |
| AGAP027997 | LRIM5 | LRIM | 1.452283 | 0.000193 | 28.893 | 0.00706 | | 1 |
| AGAP001798 | SP217 | ModSP | 1.451553 | 0.008282 | 29.136 | 0.001624 | | 1 |
| AGAP006267 | CTL6 | CTL | 1.448079 | 0.001348 | 24.568 | 0.000593 | | 1 |
| AGAP004858 | CLIPD11 | cSP_D | 1.441312 | 0 | 29.6831 | 0.00825 | | 1 |
| AGAP004981 | PPO4 | PPO | 1.440711 | 0.007255 | 27.5659 | 0.001061 | | 0 |
| AGAP010833 | CLIPB14 | cSP_B | 1.386174 | 0 | 29.1515 | 0.00888 | | 1 |
| AGAP009251 | CLIPE16 | cSPH_E | 1.114766 | 0 | 26.8062 | 0.000467 | | 2 |
| AGAP028102 | CLIPE34 | cSPH_E | 1.025307 | 0.011043 | 21.0302 | 0 | | 0 |
| AGAP028641 | CLIPE29 | cSPH_E | 1.018378 | 0.011043 | 25.9377 | 0.000404 | | 2 |
| AGAP003691 | CLIPE12 | cSPH_E | 1.017241 | 0 | 25.3616 | 0.000393 | | 2 |
| AGAP007411 | CTLMA1 | CTL | 0.972903 | 0.011043 | 22.1344 | 0.000484 | | 1 |
| AGAP003245 | CLIPA19 | cSPH_A | 0.964442 | 0.012583 | 30.2806 | 0.003796 | | 0 |
| AGAP008835 | CLIPC1 | cSP_C | 0.963808 | 0.000321 | 23.6812 | 0.000002 | | 0 |
| AGAP008366 | TEP2 | TEP | 0.960433 | 0 | 23.46 | 0.000004 | | 0 |
| AGAP009220 | CLIPB44 | cSP_B | 0.953443 | 0 | 25.3352 | 0.000097 | | 0 |
| AGAP003249 | CLIPB3b | cSP_B | 0.948186 | 0 | 28.7585 | 0.004555 | | 1 |
| AGAP003139 | SRPN9 | SRPN | 0.638504 | 0 | 18.9252 | 0.00001 | | 1 |
| AGAP007691 | SRPN18 | SRPN | 0.539241 | 0 | 23.3052 | 0.000099 | | 1 |
| AGAP007035 | APL1B | LRIM | 0.530746 | 0 | 0.53075 | 0 | | 4 |
| AGAP007036 | APL1A | LRIM | 0.530746 | 0 | 0.53075 | 0 | | 4 |
| AGAP006327 | LRIM6 | LRIM | 0.530686 | 0 | 0.53069 | 0 | | 3 |
| AGAP006416 | SP24D | SP | 0.530686 | 0 | 0.53069 | 0 | | 3 |
| AGAP028075 | CLIPE9 | cSP_E | 0.522446 | 0 | 16.9308 | 0 | | 0 |
| AGAP007412 | CTLMA3 | CTL | 0.510299 | 0 | 17.5194 | 0.000014 | | 1 |
| AGAP008403 | CLIPE15 | cSP_E | 0.5083 | 0 | 24.3571 | 0.000648 | | 1 |
| AGAP028229 | CLIPE28 | cSPH_E | 0.505625 | 0 | 20.7075 | 0 | | 0 |
| AGAP028010 | CLIPE11 | cSP_E | 0.503081 | 0 | 19.8793 | 0.000012 | | 2 |
| AGAP005496 | LRIM12 | LRIM | 0.500578 | 0 | 23.3928 | 0.000007 | | 0 |
| AGAP008091 | CLIPE1 | cSPH_E | 0.482061 | 0 | 23.6863 | 0.000321 | | 1 |
| AGAP012504 | CLIPE33 | cSP_E | 0.481607 | 0 | 24.1701 | 0.001516 | | 1 |
| AGAP010196 | CTLGA1 | CTL | 0.447654 | 0 | 21.5243 | 0.000174 | | 1 |

### Online suppl. Table S3. Undirected edge list and edge weight of AgMelGCN

| Undirected Edge | | Edge | Undirected Edge | | Edge | Undirected Edge | | Edge | Undirected Edge | | Edge Weigh |
|---|---|---|---|---|---|---|---|---|---|---|---|
| Node 1 | Node 2 | Weight | Node 1 | Node 2 | Weight | Node 1 | Node 2 | Weight | Node 1 | Node 2 | |
| PPO1 | PPO3 | 0.587719 | PPO5 | CLIPE10 | 0.567817 | PPO8 | CLIPE25 | 0.511658 | TEP3 | CLIPA5 | 0.529271 |
| PPO1 | PPO7 | 0.501504 | PPO5 | CLIPE17 | 0.475384 | PPO9 | LRIM8A | 0.592723 | TEP3 | CLIPA7 | 0.464419 |
| PPO1 | SRPN12 | 0.551528 | PPO5 | SP213 | 0.504499 | PPO9 | LRIM10 | 0.521156 | TEP3 | CLIPA8 | 0.606837 |
| PPO1 | SRPN19 | 0.507456 | PPO6 | PPO9 | 0.463975 | PPO9 | LRIM8B | 0.498801 | TEP3 | CLIPA9 | 0.515134 |
| PPO1 | CTLMA6 | 0.576697 | PPO6 | TEP1 | 0.481077 | TEP4 | TEP3 | 0.558397 | TEP3 | CLIPB10 | 0.48745 |
| PPO1 | CLIPA10 | 0.592518 | PPO6 | TEP3 | 0.564686 | TEP4 | TEP12 | 0.497159 | TEP3 | CLIPB13 | 0.487056 |
| PPO1 | CLIPA15 | 0.604387 | PPO6 | TEP15 | 0.522426 | TEP4 | APL1C | 0.526746 | TEP3 | CLIPB15 | 0.546339 |
| PPO1 | CLIPB41 | 0.445372 | PPO6 | TEP12 | 0.498961 | TEP4 | LRIM3 | 0.445251 | TEP3 | CLIPB17 | 0.560747 |
| PPO1 | CLIPD2 | 0.538257 | PPO6 | SRPN11 | 0.492822 | TEP4 | SPCLIP1 | 0.469057 | TEP3 | CLIPB2 | 0.515952 |
| PPO1 | CLIPD3 | 0.444445 | PPO6 | SRPN3 | 0.497279 | TEP4 | CLIPA5 | 0.52208 | TEP3 | CLIPB20 | 0.573417 |
| PPO1 | CLIPD7 | 0.440649 | PPO6 | SRPN5 | 0.44359 | TEP4 | CLIPA7 | 0.45957 | TEP3 | CLIPC12 | 0.642751 |
| PPO1 | CLIPD12 | 0.599786 | PPO6 | LRIM1 | 0.551392 | TEP4 | CLIPA8 | 0.450631 | TEP3 | CLIPC7 | 0.502331 |
| PPO1 | CLIPD13 | 0.487675 | PPO6 | APL1C | 0.531476 | TEP4 | CLIPB12 | 0.490684 | TEP3 | CLIPC9 | 0.465012 |
| PPO1 | CLIPD22 | 0.678472 | PPO6 | LRIM4 | 0.569444 | TEP4 | CLIPB46 | 0.487565 | TEP3 | CLIPD1 | 0.462345 |
| PPO1 | CLIPE23 | 0.443061 | PPO6 | LRIM7 | 0.444719 | TEP4 | CLIPC7 | 0.493408 | TEP3 | CLIPD6 | 0.476798 |
| PPO2 | PPO3 | 0.573265 | PPO6 | CTL4 | 0.499739 | TEP6 | TEP11 | 0.64869 | TEP3 | CLIPE10 | 0.491005 |
| PPO2 | PPO4 | 0.524013 | PPO6 | CTLMA2 | 0.511575 | TEP6 | TEP10 | 0.610704 | TEP3 | SP213 | 0.582005 |
| PPO2 | SRPN7 | 0.498167 | PPO6 | CLIPA1 | 0.553828 | TEP6 | TEP9 | 0.728272 | TEP11 | TEP10 | 0.761495 |
| PPO3 | PPO4 | 0.451847 | PPO6 | CLIPA2 | 0.499701 | TEP6 | CTL7 | 0.542676 | TEP11 | TEP9 | 0.735853 |
| PPO3 | PPO7 | 0.49531 | PPO6 | CLIPA27 | 0.606218 | TEP6 | CTLMA7 | 0.581392 | TEP11 | LRIM11 | 0.522312 |
| PPO3 | SRPN12 | 0.569369 | PPO6 | SPCLIP1 | 0.448848 | TEP1 | TEP3 | 0.561612 | TEP11 | CTL10 | 0.468067 |
| PPO3 | SRPN7 | 0.49239 | PPO6 | CLIPA4 | 0.487653 | TEP1 | SRPN14 | 0.598553 | TEP11 | CTL7 | 0.447035 |
| PPO3 | CTLGA2 | 0.479934 | PPO6 | CLIPA7 | 0.558364 | TEP1 | SRPN5 | 0.515659 | TEP11 | CTLMA7 | 0.623528 |
| PPO3 | CTLMA6 | 0.446878 | PPO6 | CLIPA8 | 0.602867 | TEP1 | LRIM1 | 0.513587 | TEP11 | CLIPB42 | 0.514947 |
| PPO3 | CLIPA10 | 0.454679 | PPO6 | CLIPA9 | 0.591665 | TEP1 | LRIM4 | 0.513977 | TEP11 | CLIPD3 | 0.526105 |
| PPO3 | CLIPD2 | 0.527588 | PPO6 | CLIPB1 | 0.519075 | TEP1 | CTL4 | 0.519652 | TEP11 | CLIPE13 | 0.461929 |
| PPO3 | CLIPD22 | 0.502254 | PPO6 | CLIPB10 | 0.529142 | TEP1 | CTLMA2 | 0.589005 | TEP11 | CLIPE30 | 0.614771 |
| PPO4 | CLIPD1 | 0.464851 | PPO6 | CLIPB13 | 0.616844 | TEP1 | CLIPA27 | 0.446179 | TEP11 | SP212 | 0.527067 |
| PPO5 | PPO6 | 0.68761 | PPO6 | CLIPB15 | 0.582959 | TEP1 | CLIPA8 | 0.615912 | TEP10 | TEP9 | 0.776146 |
| PPO5 | PPO9 | 0.462594 | PPO6 | CLIPB2 | 0.562552 | TEP1 | CLIPA9 | 0.45402 | TEP10 | TEP19 | 0.521386 |
| PPO5 | TEP1 | 0.507266 | PPO6 | CLIPB4 | 0.621076 | TEP1 | CLIPB10 | 0.55075 | TEP10 | CTL10 | 0.531597 |
| PPO5 | TEP3 | 0.490259 | PPO6 | CLIPB7 | 0.442136 | TEP1 | CLIPB13 | 0.505195 | TEP10 | CTLMA7 | 0.608985 |
| PPO5 | TEP15 | 0.446339 | PPO6 | CLIPB8 | 0.485891 | TEP1 | CLIPB15 | 0.583452 | TEP10 | CLIPA31 | 0.441405 |
| PPO5 | SRPN14 | 0.507668 | PPO6 | CLIPC12 | 0.466874 | TEP1 | CLIPB17 | 0.485988 | TEP10 | CLIPB42 | 0.485098 |
| PPO5 | SRPN3 | 0.447512 | PPO6 | CLIPC3 | 0.591954 | TEP1 | CLIPB19 | 0.56058 | TEP10 | CLIPC14 | 0.506728 |
| PPO5 | SRPN5 | 0.515494 | PPO6 | CLIPC7 | 0.470828 | TEP1 | CLIPB2 | 0.509268 | TEP10 | CLIPE13 | 0.446505 |
| PPO5 | APL1C | 0.451595 | PPO6 | CLIPC9 | 0.563786 | TEP1 | CLIPB20 | 0.487286 | TEP10 | CLIPE30 | 0.509595 |
| PPO5 | LRIM4 | 0.549572 | PPO6 | CLIPD6 | 0.552006 | TEP1 | CLIPB3a | 0.534023 | TEP10 | SP212 | 0.516337 |
| PPO5 | CTL4 | 0.473978 | PPO6 | CLIPE4 | 0.496691 | TEP1 | CLIPB4 | 0.439792 | TEP9 | CTL10 | 0.531154 |
| PPO5 | CTLMA2 | 0.458716 | PPO6 | CLIPE5 | 0.576964 | TEP1 | CLIPC12 | 0.505563 | TEP9 | CTL9 | 0.459358 |
| PPO5 | CLIPA1 | 0.452025 | PPO6 | CLIPE10 | 0.510382 | TEP1 | CLIPC4 | 0.447071 | TEP9 | CTLMA5 | 0.461774 |
| PPO5 | CLIPA2 | 0.476393 | PPO6 | SP213 | 0.521935 | TEP1 | CLIPD1 | 0.449008 | TEP9 | CTLMA7 | 0.548201 |
| PPO5 | CLIPA27 | 0.44165 | PPO7 | TEP8 | 0.494342 | TEP1 | CLIPE5 | 0.439902 | TEP9 | CLIPA31 | 0.502599 |
| PPO5 | SPCLIP1 | 0.449379 | PPO7 | CTL10 | 0.528226 | TEP1 | CLIPE6 | 0.445229 | TEP9 | CLIPB42 | 0.477594 |
| PPO5 | CLIPA8 | 0.545046 | PPO7 | CTLMA6 | 0.517259 | TEP1 | SP213 | 0.503648 | TEP9 | CLIPB45 | 0.512478 |
| PPO5 | CLIPA9 | 0.619806 | PPO7 | CTLSE1 | 0.495277 | TEP3 | TEP12 | 0.606833 | TEP9 | CLIPC14 | 0.453121 |
| PPO5 | CLIPB10 | 0.518739 | PPO7 | CLIPA10 | 0.522633 | TEP3 | SRPN1 | 0.450248 | TEP9 | CLIPD3 | 0.446627 |
| PPO5 | CLIPB13 | 0.49305 | PPO7 | CLIPA15 | 0.49713 | TEP3 | SRPN3 | 0.477913 | TEP9 | CLIPE13 | 0.494203 |
| PPO5 | CLIPB15 | 0.491874 | PPO7 | CLIPB3a | 0.497324 | TEP3 | LRIM1 | 0.472948 | TEP9 | CLIPE17 | 0.503713 |
| PPO5 | CLIPB19 | 0.502002 | PPO7 | CLIPD12 | 0.496891 | TEP3 | APL1C | 0.659159 | TEP9 | SP212 | 0.578429 |
| PPO5 | CLIPB2 | 0.583347 | PPO7 | CLIPD13 | 0.507779 | TEP3 | LRIM4 | 0.465527 | TEP8 | TEP19 | 0.717694 |
| PPO5 | CLIPB4 | 0.58336 | PPO7 | CLIPE25 | 0.515338 | TEP3 | CTL4 | 0.576196 | TEP8 | CTL9 | 0.555463 |
| PPO5 | CLIPC12 | 0.536704 | PPO8 | TEP8 | 0.487523 | TEP3 | CTLMA2 | 0.640475 | TEP8 | CLIPB42 | 0.550083 |
| PPO5 | CLIPC3 | 0.547982 | PPO8 | SRPN19 | 0.573599 | TEP3 | CLIPA1 | 0.555206 | TEP8 | CLIPD20 | 0.546118 |
| PPO5 | CLIPC9 | 0.456881 | PPO8 | CTL3 | 0.482391 | TEP3 | CLIPA2 | 0.486594 | TEP8 | CLIPE23 | 0.484535 |
| PPO5 | CLIPD6 | 0.449272 | PPO8 | CLIPB41 | 0.470245 | TEP3 | CLIPA26 | 0.489333 | TEP19 | TEP2 | 0.495861 |
| PPO5 | CLIPE4 | 0.567733 | PPO8 | CLIPB45 | 0.464128 | TEP3 | CLIPA27 | 0.547535 | TEP19 | CLIPA19 | 0.478864 |
| PPO5 | CLIPE5 | 0.676534 | PPO8 | CLIPD4 | 0.47661 | TEP3 | SPCLIP1 | 0.549916 | TEP19 | CLIPB42 | 0.579197 |
| PPO5 | CLIPE6 | 0.51216 | PPO8 | CLIPD22 | 0.484384 | TEP3 | CLIPA4 | 0.472162 | TEP19 | CLIPD20 | 0.521403 |

| Undirected Edge | | Edge Weight | Undirected Edge | | Edge Weight | Undirected Edge | | Edge Weight | Undirected Edge | | Edge Weigh |
|---|---|---|---|---|---|---|---|---|---|---|---|
| Node 1 | Node 2 | | Node 1 | Node 2 | | Node 1 | Node 2 | | Node 1 | Node 2 | |
| TEP15 | SRPN11 | 0.480188 | TEP12 | CLIPA7 | 0.494523 | SRPN12 | CLIPD13 | 0.480268 | SRPN2 | CLIPB8 | 0.503529 |
| TEP15 | SRPN14 | 0.465307 | TEP12 | CLIPA8 | 0.651934 | SRPN12 | CLIPD14 | 0.470523 | SRPN2 | CLIPC3 | 0.533303 |
| TEP15 | SRPN2 | 0.457021 | TEP12 | CLIPA9 | 0.51413 | SRPN12 | CLIPD22 | 0.536664 | SRPN2 | SP213 | 0.466565 |
| TEP15 | SRPN3 | 0.557435 | TEP12 | CLIPB1 | 0.439384 | SRPN14 | SRPN17 | 0.462225 | SRPN3 | SRPN4 | 0.493668 |
| TEP15 | LRIM1 | 0.463721 | TEP12 | CLIPB12 | 0.481061 | SRPN14 | SRPN5 | 0.489598 | SRPN3 | LRIM1 | 0.538519 |
| TEP15 | APL1C | 0.514173 | TEP12 | CLIPB15 | 0.517046 | SRPN14 | CLIPA4 | 0.476267 | SRPN3 | APL1C | 0.654259 |
| TEP15 | LRIM4 | 0.45289 | TEP12 | CLIPB2 | 0.600329 | SRPN14 | CLIPA9 | 0.442825 | SRPN3 | LRIM4 | 0.5492 |
| TEP15 | CTLGA3 | 0.439662 | TEP12 | CLIPB20 | 0.493418 | SRPN14 | CLIPB10 | 0.467426 | SRPN3 | CTL4 | 0.477549 |
| TEP15 | CLIPA2 | 0.512554 | TEP12 | CLIPB8 | 0.527456 | SRPN14 | CLIPB11 | 0.499532 | SRPN3 | CTLMA2 | 0.463205 |
| TEP15 | CLIPA4 | 0.491949 | TEP12 | CLIPB46 | 0.535876 | SRPN14 | CLIPB4 | 0.588831 | SRPN3 | CLIPA1 | 0.4763 |
| TEP15 | CLIPA7 | 0.640558 | TEP12 | CLIPC12 | 0.597669 | SRPN14 | CLIPB47 | 0.47821 | SRPN3 | CLIPA2 | 0.607445 |
| TEP15 | CLIPA8 | 0.526491 | TEP12 | CLIPC3 | 0.451921 | SRPN14 | CLIPC12 | 0.504719 | SRPN3 | CLIPA26 | 0.439178 |
| TEP15 | CLIPA9 | 0.519561 | TEP12 | CLIPC7 | 0.509635 | SRPN14 | CLIPC3 | 0.510417 | SRPN3 | CLIPA27 | 0.587976 |
| TEP15 | CLIPB1 | 0.497962 | TEP12 | CLIPC9 | 0.454448 | SRPN14 | CLIPC4 | 0.614795 | SRPN3 | CLIPA4 | 0.46589 |
| TEP15 | CLIPB10 | 0.480182 | TEP12 | CLIPE4 | 0.460732 | SRPN14 | SP213 | 0.446599 | SRPN3 | CLIPA7 | 0.639114 |
| TEP15 | CLIPB13 | 0.509055 | TEP12 | CLIPE5 | 0.500204 | SRPN16 | SRPN3 | 0.475264 | SRPN3 | CLIPA8 | 0.620381 |
| TEP15 | CLIPB4 | 0.60236 | TEP12 | CLIPE10 | 0.547677 | SRPN16 | CTLMA8 | 0.48981 | SRPN3 | CLIPA9 | 0.562592 |
| TEP15 | CLIPB8 | 0.531852 | SRPN_x | SRPN6 | 0.479074 | SRPN16 | CLIPA7 | 0.455619 | SRPN3 | CLIPB10 | 0.477085 |
| TEP15 | CLIPC3 | 0.511804 | SRPN_x | SRPN9 | 0.638504 | SRPN16 | CLIPB11 | 0.678085 | SRPN3 | CLIPB13 | 0.503884 |
| TEP15 | CLIPC6 | 0.487043 | SRPN_x | CTL6 | 0.480687 | SRPN16 | CLIPB4 | 0.454544 | SRPN3 | CLIPB15 | 0.560841 |
| TEP15 | CLIPC7 | 0.572819 | SRPN1 | SRPN3 | 0.496837 | SRPN16 | CLIPB5 | 0.479272 | SRPN3 | CLIPB2 | 0.453117 |
| TEP15 | CLIPC9 | 0.459722 | SRPN1 | SRPN4 | 0.541593 | SRPN17 | SRPN3 | 0.460616 | SRPN3 | CLIPB4 | 0.504196 |
| TEP15 | CLIPD6 | 0.493569 | SRPN1 | APL1C | 0.507615 | SRPN17 | SRPN5 | 0.548226 | SRPN3 | CLIPB8 | 0.499749 |
| TEP15 | CLIPE4 | 0.542196 | SRPN1 | CLIPA27 | 0.494203 | SRPN17 | SRPN6 | 0.442257 | SRPN3 | CLIPC7 | 0.47096 |
| TEP15 | SP213 | 0.445078 | SRPN1 | CLIPA8 | 0.479238 | SRPN17 | SRPN8 | 0.462298 | SRPN3 | CLIPC9 | 0.536706 |
| TEP2 | CLIPB42 | 0.464572 | SRPN11 | SRPN2 | 0.638378 | SRPN17 | CTLMA2 | 0.504346 | SRPN3 | SP213 | 0.507729 |
| TEP14 | TEP12 | 0.618507 | SRPN11 | SRPN4 | 0.465264 | SRPN17 | CLIPA7 | 0.503175 | SRPN4 | APL1C | 0.467324 |
| TEP14 | APL1C | 0.582411 | SRPN11 | LRIM1 | 0.625216 | SRPN17 | CLIPB10 | 0.517169 | SRPN4 | CTLMA1 | 0.462603 |
| TEP14 | LRIM3 | 0.661812 | SRPN11 | APL1C | 0.538737 | SRPN17 | CLIPB11 | 0.441608 | SRPN4 | CLIPA7 | 0.453537 |
| TEP14 | LRIM15 | 0.53547 | SRPN11 | CTL4 | 0.460314 | SRPN17 | CLIPB16 | 0.463485 | SRPN5 | SRPN6 | 0.54365 |
| TEP14 | LRIM9 | 0.484625 | SRPN11 | CTLMA2 | 0.495158 | SRPN17 | CLIPB2 | 0.463908 | SRPN5 | LRIM1 | 0.439892 |
| TEP14 | CLIPA2 | 0.550799 | SRPN11 | CLIPA1 | 0.580122 | SRPN17 | CLIPB4 | 0.515167 | SRPN5 | LRIM4 | 0.480886 |
| TEP14 | CLIPA26 | 0.581263 | SRPN11 | CLIPA14 | 0.601313 | SRPN17 | CLIPB7 | 0.528401 | SRPN5 | CTL4 | 0.461222 |
| TEP14 | CLIPA27 | 0.529932 | SRPN11 | CLIPA27 | 0.459709 | SRPN17 | CLIPC3 | 0.445177 | SRPN5 | CTLMA2 | 0.647642 |
| TEP14 | CLIPA5 | 0.501293 | SRPN11 | SPCLIP1 | 0.496314 | SRPN17 | CLIPD11 | 0.459407 | SRPN5 | CLIPA7 | 0.551312 |
| TEP14 | CLIPA8 | 0.468549 | SRPN11 | CLIPA4 | 0.59774 | SRPN18 | CLIPB47 | 0.539241 | SRPN5 | CLIPA9 | 0.502432 |
| TEP14 | CLIPB12 | 0.513375 | SRPN11 | CLIPA6 | 0.573996 | SRPN19 | CTLMA7 | 0.562293 | SRPN5 | CLIPB10 | 0.471721 |
| TEP14 | CLIPB2 | 0.499803 | SRPN11 | CLIPA7 | 0.63922 | SRPN19 | CTLMA9 | 0.465524 | SRPN5 | CLIPB16 | 0.448297 |
| TEP14 | CLIPB46 | 0.551365 | SRPN11 | CLIPA8 | 0.616997 | SRPN19 | CLIPB41 | 0.468072 | SRPN5 | CLIPB4 | 0.469149 |
| TEP14 | CLIPE4 | 0.474565 | SRPN11 | CLIPA9 | 0.682315 | SRPN19 | CLIPC13 | 0.457837 | SRPN5 | CLIPB7 | 0.517617 |
| TEP14 | CLIPE5 | 0.457386 | SRPN11 | CLIPB1 | 0.556095 | SRPN19 | CLIPD12 | 0.4535 | SRPN5 | CLIPC3 | 0.55008 |
| TEP14 | CLIPE10 | 0.551857 | SRPN11 | CLIPB10 | 0.497292 | SRPN19 | CLIPD22 | 0.467528 | SRPN5 | CLIPD6 | 0.454406 |
| TEP13 | SRPN12 | 0.48343 | SRPN11 | CLIPB13 | 0.64641 | SRPN19 | CLIPE13 | 0.446727 | SRPN5 | CLIPD11 | 0.457057 |
| TEP13 | SRPN7 | 0.45757 | SRPN11 | CLIPB15 | 0.536253 | SRPN19 | CLIPE24 | 0.483996 | SRPN5 | CLIPE4 | 0.531239 |
| TEP13 | CLIPD22 | 0.52124 | SRPN11 | CLIPB2 | 0.455831 | SRPN19 | CLIPE27 | 0.531675 | SRPN5 | CLIPE5 | 0.466613 |
| TEP12 | SRPN11 | 0.488455 | SRPN11 | CLIPB4 | 0.53485 | SRPN19 | CLIPE30 | 0.49282 | SRPN5 | CLIPE6 | 0.614617 |
| TEP12 | SRPN3 | 0.456032 | SRPN11 | CLIPB5 | 0.488816 | SRPN2 | LRIM1 | 0.54414 | SRPN5 | SP213 | 0.684213 |
| TEP12 | LRIM1 | 0.51081 | SRPN11 | CLIPB8 | 0.541947 | SRPN2 | CTLMA2 | 0.440775 | SRPN6 | CTL6 | 0.45125 |
| TEP12 | APL1C | 0.664364 | SRPN11 | CLIPC3 | 0.630281 | SRPN2 | CLIPA1 | 0.487439 | SRPN6 | CLIPA7 | 0.492704 |
| TEP12 | LRIM3 | 0.591741 | SRPN11 | CLIPC7 | 0.462124 | SRPN2 | CLIPA14 | 0.598725 | SRPN7 | SRPN8 | 0.509969 |
| TEP12 | LRIM4 | 0.456708 | SRPN11 | CLIPC9 | 0.515256 | SRPN2 | CLIPA4 | 0.533058 | SRPN7 | CTLGA2 | 0.492249 |
| TEP12 | LRIM9 | 0.453299 | SRPN11 | CLIPE5 | 0.448418 | SRPN2 | CLIPA6 | 0.601288 | SRPN7 | CTLMA8 | 0.501003 |
| TEP12 | CTL4 | 0.539719 | SRPN11 | SP213 | 0.603114 | SRPN2 | CLIPA7 | 0.572744 | SRPN7 | CLIPA10 | 0.467974 |
| TEP12 | CTLMA2 | 0.463289 | SRPN12 | CTLGA2 | 0.512773 | SRPN2 | CLIPA8 | 0.444925 | SRPN7 | CLIPD2 | 0.493592 |
| TEP12 | CLIPA1 | 0.572956 | SRPN12 | CTLMA6 | 0.493594 | SRPN2 | CLIPA9 | 0.522495 | SRPN7 | CLIPD7 | 0.45081 |
| TEP12 | CLIPA2 | 0.53193 | SRPN12 | CLIPA10 | 0.44933 | SRPN2 | CLIPB1 | 0.558203 | SRPN8 | CLIPA4 | 0.461107 |
| TEP12 | CLIPA26 | 0.546017 | SRPN12 | CLIPD2 | 0.501755 | SRPN2 | CLIPB13 | 0.612758 | SRPN8 | CLIPB11 | 0.474369 |
| TEP12 | CLIPA27 | 0.592893 | SRPN12 | CLIPD8 | 0.445837 | SRPN2 | CLIPB15 | 0.455761 | LRIM5 | LRIM7 | 0.471828 |
| TEP12 | CLIPA5 | 0.589518 | SRPN12 | CLIPD12 | 0.454167 | SRPN2 | CLIPB4 | 0.633261 | LRIM5 | CLIPA2 | 0.493836 |

38

| Undirected Edge | | Edge | Undirected Edge | | Edge | Undirected Edge | | Edge | Undirected Edge | | Edge Weigh |
|---|---|---|---|---|---|---|---|---|---|---|---|
| Node 1 | Node 2 | Weight | Node 1 | Node 2 | Weight | Node 1 | Node 2 | Weight | Node 1 | Node 2 | |
| LRIM5 | CLIPA26 | 0.486619 | APL1C | CLIPC12 | 0.505165 | LRIM7 | CLIPB12 | 0.512666 | CTL9 | CLIPB42 | 0.619[illegible] |
| LRIM12 | CLIPC13 | 0.500578 | APL1C | CLIPC7 | 0.589653 | LRIM7 | CLIPB2 | 0.476882 | CTL9 | CLIPC14 | 0.54582[illegible] |
| LRIM6 | SP24D | 0.530686 | APL1C | CLIPC9 | 0.526246 | LRIM7 | CLIPB4 | 0.444096 | CTL9 | CLIPD3 | 0.50677[illegible] |
| LRIM1 | APL1C | 0.63015 | APL1C | CLIPE4 | 0.589585 | LRIM7 | CLIPC6 | 0.502384 | CTL9 | CLIPD4 | 0.47285[illegible] |
| LRIM1 | CTL4 | 0.620526 | APL1C | CLIPE5 | 0.57524 | LRIM7 | CLIPE5 | 0.460276 | CTL9 | CLIPD8 | 0.47016[illegible] |
| LRIM1 | CTLGA3 | 0.467321 | APL1C | CLIPE10 | 0.531791 | CTL1 | CTL3 | 0.508238 | CTL9 | CLIPE25 | 0.52109[illegible] |
| LRIM1 | CTLMA2 | 0.698006 | APL1C | SP213 | 0.586385 | CTL1 | CLIPD4 | 0.469561 | CTLGA1 | CLIPB9 | 0.44765[illegible] |
| LRIM1 | CLIPA1 | 0.744573 | LRIM11 | CLIPC13 | 0.555437 | CTL1 | CLIPE25 | 0.532374 | CTLGA2 | CTLMA6 | 0.46940[illegible] |
| LRIM1 | CLIPA14 | 0.541812 | LRIM11 | CLIPE2 | 0.476745 | CTL10 | CTL7 | 0.62812 | CTLGA2 | CLIPD8 | 0.48353[illegible] |
| LRIM1 | CLIPA2 | 0.62029 | LRIM11 | SP214 | 0.49593 | CTL10 | CTLMA7 | 0.56964 | CTLGA2 | CLIPE25 | 0.52186[illegible] |
| LRIM1 | CLIPA27 | 0.539769 | APL1B | APL1A | 0.530746 | CTL10 | CTLMA9 | 0.45779 | CTLGA3 | CLIPA6 | 0.48913[illegible] |
| LRIM1 | SPCLIP1 | 0.648172 | LRIM3 | LRIM9 | 0.472662 | CTL10 | CLIPA15 | 0.489542 | CTLGA3 | CLIPB4 | 0.44471[illegible] |
| LRIM1 | CLIPA4 | 0.460986 | LRIM3 | CLIPA1 | 0.459254 | CTL10 | CLIPA31 | 0.540756 | CTLGA3 | CLIPD6 | 0.49052[illegible] |
| LRIM1 | CLIPA6 | 0.52612 | LRIM3 | CLIPA2 | 0.465525 | CTL10 | CLIPA32 | 0.489134 | CTLMA1 | CTLMA3 | 0.51029[illegible] |
| LRIM1 | CLIPA7 | 0.658819 | LRIM3 | CLIPA26 | 0.456378 | CTL10 | CLIPD3 | 0.447673 | CTLMA2 | CLIPA1 | 0.71707[illegible] |
| LRIM1 | CLIPA8 | 0.644312 | LRIM3 | CLIPA27 | 0.557849 | CTL10 | CLIPD12 | 0.507072 | CTLMA2 | CLIPA14 | 0.50621[illegible] |
| LRIM1 | CLIPA9 | 0.729312 | LRIM3 | CLIPB2 | 0.462548 | CTL10 | CLIPE13 | 0.448296 | CTLMA2 | CLIPA2 | 0.50636[illegible] |
| LRIM1 | CLIPB1 | 0.465749 | LRIM3 | CLIPB46 | 0.520941 | CTL10 | CLIPE18 | 0.444632 | CTLMA2 | CLIPA27 | 0.59604[illegible] |
| LRIM1 | CLIPB10 | 0.562118 | LRIM3 | CLIPC7 | 0.477494 | CTL10 | CLIPE21 | 0.590841 | CTLMA2 | SPCLIP1 | 0.71646[illegible] |
| LRIM1 | CLIPB12 | 0.481919 | LRIM4 | CTL4 | 0.466319 | CTL10 | CLIPE25 | 0.533543 | CTLMA2 | CLIPA4 | 0.46801[illegible] |
| LRIM1 | CLIPB13 | 0.575467 | LRIM4 | CTLMA2 | 0.459957 | CTL10 | SP212 | 0.448822 | CTLMA2 | CLIPA7 | 0.58485[illegible] |
| LRIM1 | CLIPB15 | 0.667307 | LRIM4 | CLIPA1 | 0.457147 | CTL3 | CLIPE25 | 0.525951 | CTLMA2 | CLIPA8 | 0.69410[illegible] |
| LRIM1 | CLIPB17 | 0.470871 | LRIM4 | CLIPA2 | 0.518855 | CTL4 | CTLMA2 | 0.781166 | CTLMA2 | CLIPA9 | 0.74083[illegible] |
| LRIM1 | CLIPB2 | 0.508549 | LRIM4 | CLIPA27 | 0.472369 | CTL4 | CLIPA1 | 0.670683 | CTLMA2 | CLIPB10 | 0.62361[illegible] |
| LRIM1 | CLIPB4 | 0.528836 | LRIM4 | CLIPA7 | 0.506429 | CTL4 | CLIPA2 | 0.591561 | CTLMA2 | CLIPB13 | 0.50641[illegible] |
| LRIM1 | CLIPB7 | 0.444594 | LRIM4 | CLIPA8 | 0.573786 | CTL4 | CLIPA27 | 0.556836 | CTLMA2 | CLIPB15 | 0.71720[illegible] |
| LRIM1 | CLIPB8 | 0.524387 | LRIM4 | CLIPA9 | 0.608276 | CTL4 | SPCLIP1 | 0.534676 | CTLMA2 | CLIPB2 | 0.54947[illegible] |
| LRIM1 | CLIPC12 | 0.542033 | LRIM4 | CLIPB1 | 0.441563 | CTL4 | CLIPA7 | 0.515505 | CTLMA2 | CLIPB20 | 0.52550[illegible] |
| LRIM1 | CLIPC3 | 0.491121 | LRIM4 | CLIPB10 | 0.544269 | CTL4 | CLIPA8 | 0.709674 | CTLMA2 | CLIPB4 | 0.51770[illegible] |
| LRIM1 | CLIPC7 | 0.537464 | LRIM4 | CLIPB15 | 0.486781 | CTL4 | CLIPA9 | 0.721854 | CTLMA2 | CLIPB7 | 0.62135[illegible] |
| LRIM1 | CLIPC9 | 0.511065 | LRIM4 | CLIPB2 | 0.511386 | CTL4 | CLIPB10 | 0.528875 | CTLMA2 | CLIPB8 | 0.44422[illegible] |
| LRIM1 | CLIPD6 | 0.50167 | LRIM4 | CLIPB20 | 0.442573 | CTL4 | CLIPB12 | 0.460007 | CTLMA2 | CLIPC12 | 0.61157[illegible] |
| LRIM1 | CLIPE4 | 0.473958 | LRIM4 | CLIPB4 | 0.515524 | CTL4 | CLIPB13 | 0.446543 | CTLMA2 | CLIPC3 | 0.61459[illegible] |
| LRIM1 | CLIPE5 | 0.558266 | LRIM4 | CLIPB7 | 0.455762 | CTL4 | CLIPB15 | 0.686282 | CTLMA2 | CLIPC9 | 0.55381[illegible] |
| LRIM1 | SP213 | 0.611351 | LRIM4 | CLIPC12 | 0.533916 | CTL4 | CLIPB19 | 0.470962 | CTLMA2 | CLIPD1 | 0.47090[illegible] |
| APL1C | LRIM3 | 0.518785 | LRIM4 | CLIPC3 | 0.447797 | CTL4 | CLIPB2 | 0.548608 | CTLMA2 | CLIPD6 | 0.53708[illegible] |
| APL1C | LRIM4 | 0.517662 | LRIM4 | CLIPD6 | 0.458825 | CTL4 | CLIPB20 | 0.495406 | CTLMA2 | CLIPE4 | 0.54301[illegible] |
| APL1C | LRIM9 | 0.468133 | LRIM4 | CLIPE4 | 0.501482 | CTL4 | CLIPB4 | 0.466889 | CTLMA2 | CLIPE5 | 0.54340[illegible] |
| APL1C | CTL4 | 0.657798 | LRIM4 | CLIPE5 | 0.558026 | CTL4 | CLIPB7 | 0.46422 | CTLMA2 | CLIPE6 | 0.572[illegible] |
| APL1C | CTLMA2 | 0.611476 | LRIM15 | CLIPA27 | 0.543182 | CTL4 | CLIPC12 | 0.597918 | CTLMA2 | SP213 | 0.71856[illegible] |
| APL1C | CLIPA1 | 0.631048 | LRIM15 | CLIPC14 | 0.479143 | CTL4 | CLIPC3 | 0.596872 | CTLMA4 | CLIPB2 | 0.45398[illegible] |
| APL1C | CLIPA2 | 0.739474 | LRIM15 | CLIPE4 | 0.470625 | CTL4 | CLIPC9 | 0.571407 | CTLMA5 | CTLMA8 | 0.52524[illegible] |
| APL1C | CLIPA26 | 0.464671 | LRIM9 | LRIM8A | 0.712875 | CTL4 | CLIPD6 | 0.449106 | CTLMA5 | CLIPD2 | 0.51241[illegible] |
| APL1C | CLIPA27 | 0.724067 | LRIM9 | LRIM10 | 0.757774 | CTL4 | CLIPE4 | 0.501524 | CTLMA5 | CLIPD7 | 0.53343[illegible] |
| APL1C | SPCLIP1 | 0.468174 | LRIM9 | LRIM8B | 0.68824 | CTL4 | CLIPE5 | 0.588959 | CTLMA5 | SP212 | 0.46872[illegible] |
| APL1C | CLIPA5 | 0.496529 | LRIM9 | CLIPA9 | 0.524501 | CTL4 | CLIPE10 | 0.538829 | CTLMA6 | CTLSE2 | 0.44173[illegible] |
| APL1C | CLIPA7 | 0.507056 | LRIM8A | LRIM10 | 0.714055 | CTL4 | SP213 | 0.586711 | CTLMA6 | CLIPA10 | 0.68599[illegible] |
| APL1C | CLIPA8 | 0.644116 | LRIM8A | LRIM8B | 0.692411 | CTL5 | CLIPC13 | 0.542859 | CTLMA6 | CLIPA15 | 0.47204[illegible] |
| APL1C | CLIPA9 | 0.657998 | LRIM8A | CTL6 | 0.516142 | CTL5 | CLIPE2 | 0.513721 | CTLMA6 | CLIPD2 | 0.49722[illegible] |
| APL1C | CLIPB10 | 0.524308 | LRIM8A | CTLMA4 | 0.496034 | CTL5 | SP214 | 0.614979 | CTLMA6 | CLIPD3 | 0.47635[illegible] |
| APL1C | CLIPB12 | 0.534846 | LRIM8A | CLIPB2 | 0.490293 | CTL7 | CTLMA7 | 0.563022 | CTLMA6 | CLIPD7 | 0.47191[illegible] |
| APL1C | CLIPB13 | 0.555242 | LRIM8A | CLIPB3b | 0.495551 | CTL7 | CTLMA9 | 0.499793 | CTLMA6 | CLIPD12 | 0.5964[illegible] |
| APL1C | CLIPB14 | 0.458643 | LRIM8A | CLIPC3 | 0.524514 | CTL7 | CLIPA31 | 0.549003 | CTLMA6 | CLIPD13 | 0.62433[illegible] |
| APL1C | CLIPB15 | 0.65947 | LRIM10 | LRIM8B | 0.741212 | CTL7 | CLIPA32 | 0.499609 | CTLMA6 | CLIPD14 | 0.45645[illegible] |
| APL1C | CLIPB17 | 0.495379 | LRIM10 | CTLMA4 | 0.506675 | CTL7 | CLIPD12 | 0.508911 | CTLMA6 | CLIPD22 | 0.51801[illegible] |
| APL1C | CLIPB2 | 0.599427 | LRIM8B | CLIPA9 | 0.479297 | CTL7 | CLIPE2 | 0.441838 | CTLMA6 | CLIPE2 | 0.44488[illegible] |
| APL1C | CLIPB20 | 0.598461 | LRIM8B | CLIPB12 | 0.475426 | CTL7 | CLIPE21 | 0.608244 | CTLMA6 | CLIPE21 | 0.46619[illegible] |
| APL1C | CLIPB7 | 0.453737 | LRIM7 | CLIPA8 | 0.455816 | CTL7 | CLIPE22 | 0.595686 | CTLMA6 | CLIPE23 | 0.48826[illegible] |
| APL1C | CLIPB46 | 0.570886 | LRIM7 | CLIPA9 | 0.520241 | CTL9 | CLIPA31 | 0.4442 | CTLMA6 | CLIPE25 | 0.5076[illegible] |

| Undirected Edge | | Edge | Undirected Edge | | Edge | Undirected Edge | | Edge | Undirected Edge | | Edge Weigh |
|---|---|---|---|---|---|---|---|---|---|---|---|
| Node 1 | Node 2 | Weight | Node 1 | Node 2 | Weight | Node 1 | Node 2 | Weight | Node 1 | Node 2 | |
| CTLMA7 | CTLMA9 | 0.529005 | CLIPA10 | CLIPD2 | 0.516507 | CLIPA2 | CLIPC9 | 0.550117 | CLIPA4 | CLIPA6 | 0.442257 |
| CTLMA7 | CLIPA31 | 0.583517 | CLIPA10 | CLIPD3 | 0.617817 | CLIPA2 | CLIPD6 | 0.447977 | CLIPA4 | CLIPA7 | 0.611526 |
| CTLMA7 | CLIPB42 | 0.456823 | CLIPA10 | CLIPD7 | 0.672602 | CLIPA2 | CLIPE4 | 0.473877 | CLIPA4 | CLIPA9 | 0.496169 |
| CTLMA7 | CLIPD3 | 0.595815 | CLIPA10 | CLIPD8 | 0.518597 | CLIPA2 | CLIPE5 | 0.517368 | CLIPA4 | CLIPB1 | 0.504017 |
| CTLMA7 | CLIPD8 | 0.528017 | CLIPA10 | CLIPD12 | 0.714495 | CLIPA2 | CLIPE10 | 0.605037 | CLIPA4 | CLIPB10 | 0.559687 |
| CTLMA7 | CLIPD14 | 0.440476 | CLIPA10 | CLIPD13 | 0.637124 | CLIPA26 | CLIPA27 | 0.538904 | CLIPA4 | CLIPB11 | 0.650044 |
| CTLMA7 | CLIPE13 | 0.464129 | CLIPA10 | CLIPD14 | 0.480905 | CLIPA26 | CLIPA5 | 0.549758 | CLIPA4 | CLIPB13 | 0.558752 |
| CTLMA7 | CLIPE19 | 0.509271 | CLIPA10 | CLIPD22 | 0.64018 | CLIPA26 | CLIPA8 | 0.47707 | CLIPA4 | CLIPB4 | 0.622294 |
| CTLMA7 | CLIPE21 | 0.536111 | CLIPA10 | CLIPE21 | 0.472546 | CLIPA26 | CLIPB14 | 0.4458 | CLIPA4 | CLIPB5 | 0.593112 |
| CTLMA7 | SP212 | 0.446085 | CLIPA10 | CLIPE23 | 0.537944 | CLIPA26 | CLIPB15 | 0.484701 | CLIPA4 | CLIPB7 | 0.441255 |
| CTLMA8 | CLIPA10 | 0.515537 | CLIPA10 | CLIPE25 | 0.583791 | CLIPA26 | CLIPB17 | 0.462542 | CLIPA4 | CLIPB8 | 0.586303 |
| CTLMA8 | CLIPA32 | 0.556253 | CLIPA12 | CLIPA14 | 0.555334 | CLIPA26 | CLIPB2 | 0.464738 | CLIPA4 | CLIPB9 | 0.600358 |
| CTLMA9 | CLIPA31 | 0.44844 | CLIPA12 | SPCLIP1 | 0.474123 | CLIPA26 | CLIPE5 | 0.450368 | CLIPA4 | CLIPC10 | 0.538667 |
| CTLMA9 | CLIPD3 | 0.454745 | CLIPA12 | CLIPA4 | 0.582464 | CLIPA26 | CLIPE33 | 0.481607 | CLIPA4 | CLIPC3 | 0.489984 |
| CTLMA9 | CLIPE19 | 0.525174 | CLIPA12 | CLIPB1 | 0.521043 | CLIPA26 | SP217 | 0.484754 | CLIPA4 | CLIPC4 | 0.461429 |
| CTLMA9 | CLIPE21 | 0.479674 | CLIPA12 | CLIPB11 | 0.463794 | CLIPA27 | CLIPA4 | 0.455691 | CLIPA4 | CLIPC7 | 0.519369 |
| CTLMA9 | CLIPE25 | 0.494745 | CLIPA12 | CLIPB4 | 0.53439 | CLIPA27 | CLIPA5 | 0.517927 | CLIPA4 | CLIPC9 | 0.468734 |
| CTLMA9 | CLIPE27 | 0.480747 | CLIPA12 | CLIPC4 | 0.491421 | CLIPA27 | CLIPA7 | 0.651791 | CLIPA4 | CLIPD6 | 0.477692 |
| CTLSE1 | CLIPB47 | 0.446522 | CLIPA12 | CLIPC6 | 0.484134 | CLIPA27 | CLIPA8 | 0.698796 | CLIPA4 | SP213 | 0.592019 |
| CTLSE1 | CLIPD12 | 0.492842 | CLIPA13 | CLIPA14 | 0.440683 | CLIPA27 | CLIPA9 | 0.625073 | CLIPA5 | CLIPA8 | 0.469486 |
| CTLSE1 | CLIPE2 | 0.590299 | CLIPA13 | CLIPA4 | 0.534913 | CLIPA27 | CLIPB10 | 0.480225 | CLIPA5 | CLIPB12 | 0.594621 |
| CTLSE2 | CLIPA10 | 0.468365 | CLIPA13 | CLIPC13 | 0.627309 | CLIPA27 | CLIPB12 | 0.479741 | CLIPA5 | CLIPB2 | 0.488674 |
| CTLSE2 | CLIPA15 | 0.522115 | CLIPA14 | SPCLIP1 | 0.637245 | CLIPA27 | CLIPB13 | 0.481863 | CLIPA5 | CLIPB46 | 0.507914 |
| CTLSE2 | CLIPD3 | 0.572327 | CLIPA14 | CLIPA4 | 0.588574 | CLIPA27 | CLIPB15 | 0.580234 | CLIPA5 | CLIPC12 | 0.491484 |
| CTLSE2 | CLIPD8 | 0.445608 | CLIPA14 | CLIPA6 | 0.737991 | CLIPA27 | CLIPB17 | 0.580742 | CLIPA5 | CLIPC7 | 0.468546 |
| CTLSE2 | CLIPD12 | 0.439511 | CLIPA14 | CLIPA7 | 0.618924 | CLIPA27 | CLIPB2 | 0.654548 | CLIPA5 | CLIPE10 | 0.449528 |
| CTLSE2 | CLIPD13 | 0.532276 | CLIPA14 | CLIPB1 | 0.502284 | CLIPA27 | CLIPB20 | 0.490151 | CLIPA6 | CLIPA7 | 0.545222 |
| CTLSE2 | CLIPD14 | 0.515421 | CLIPA14 | CLIPB13 | 0.614034 | CLIPA27 | CLIPB3a | 0.439695 | CLIPA6 | CLIPB1 | 0.544473 |
| CLIPA1 | CLIPA14 | 0.548138 | CLIPA14 | CLIPB4 | 0.573336 | CLIPA27 | CLIPB4 | 0.482351 | CLIPA6 | CLIPB13 | 0.564449 |
| CLIPA1 | CLIPA2 | 0.658142 | CLIPA14 | CLIPB8 | 0.510802 | CLIPA27 | CLIPB8 | 0.595458 | CLIPA6 | CLIPB4 | 0.496834 |
| CLIPA1 | CLIPA27 | 0.563273 | CLIPA14 | CLIPB9 | 0.476802 | CLIPA27 | CLIPB46 | 0.564394 | CLIPA6 | CLIPB8 | 0.44034 |
| CLIPA1 | SPCLIP1 | 0.702906 | CLIPA14 | CLIPD6 | 0.4811 | CLIPA27 | CLIPC12 | 0.504624 | CLIPA6 | CLIPD6 | 0.510926 |
| CLIPA1 | CLIPA4 | 0.529153 | CLIPA14 | SP213 | 0.574606 | CLIPA27 | CLIPC3 | 0.492765 | CLIPA7 | CLIPA8 | 0.544403 |
| CLIPA1 | CLIPA5 | 0.457816 | CLIPA15 | CLIPB41 | 0.460107 | CLIPA27 | CLIPC7 | 0.471826 | CLIPA7 | CLIPA9 | 0.595802 |
| CLIPA1 | CLIPA6 | 0.539036 | CLIPA15 | CLIPD3 | 0.492944 | CLIPA27 | CLIPC9 | 0.605961 | CLIPA7 | CLIPB1 | 0.546211 |
| CLIPA1 | CLIPA7 | 0.585352 | CLIPA15 | CLIPD7 | 0.497252 | CLIPA27 | CLIPD1 | 0.471953 | CLIPA7 | CLIPB13 | 0.466452 |
| CLIPA1 | CLIPA8 | 0.659059 | CLIPA15 | CLIPD12 | 0.69611 | CLIPA27 | CLIPE4 | 0.483445 | CLIPA7 | CLIPB15 | 0.519117 |
| CLIPA1 | CLIPA9 | 0.732836 | CLIPA15 | CLIPD13 | 0.576385 | CLIPA27 | CLIPE5 | 0.500266 | CLIPA7 | CLIPB16 | 0.457305 |
| CLIPA1 | CLIPB1 | 0.501361 | CLIPA15 | CLIPD22 | 0.633821 | CLIPA27 | CLIPE10 | 0.594262 | CLIPA7 | CLIPB2 | 0.517408 |
| CLIPA1 | CLIPB10 | 0.553333 | CLIPA15 | CLIPE2 | 0.528097 | CLIPA27 | SP213 | 0.482937 | CLIPA7 | CLIPB4 | 0.650376 |
| CLIPA1 | CLIPB12 | 0.583357 | CLIPA15 | CLIPE23 | 0.623391 | CLIPA3 | CLIPD20 | 0.460774 | CLIPA7 | CLIPB5 | 0.515423 |
| CLIPA1 | CLIPB13 | 0.591383 | CLIPA15 | CLIPE25 | 0.542019 | CLIPA3 | CLIPE2 | 0.619083 | CLIPA7 | CLIPB7 | 0.546453 |
| CLIPA1 | CLIPB15 | 0.685842 | CLIPA19 | CLIPE4 | 0.485578 | CLIPA3 | CLIPE23 | 0.504084 | CLIPA7 | CLIPB8 | 0.68485 |
| CLIPA1 | CLIPB2 | 0.564269 | CLIPA2 | CLIPA26 | 0.444449 | SPCLIP1 | CLIPA4 | 0.516186 | CLIPA7 | CLIPC10 | 0.545408 |
| CLIPA1 | CLIPB20 | 0.521387 | CLIPA2 | CLIPA27 | 0.508912 | SPCLIP1 | CLIPA6 | 0.531494 | CLIPA7 | CLIPC3 | 0.565425 |
| CLIPA1 | CLIPB4 | 0.552025 | CLIPA2 | SPCLIP1 | 0.544845 | SPCLIP1 | CLIPA7 | 0.617151 | CLIPA7 | CLIPC7 | 0.472114 |
| CLIPA1 | CLIPB8 | 0.542014 | CLIPA2 | CLIPA8 | 0.689739 | SPCLIP1 | CLIPA9 | 0.554338 | CLIPA7 | CLIPC9 | 0.590511 |
| CLIPA1 | CLIPC12 | 0.657239 | CLIPA2 | CLIPA9 | 0.599436 | SPCLIP1 | CLIPB12 | 0.461774 | CLIPA7 | CLIPD6 | 0.490606 |
| CLIPA1 | CLIPC3 | 0.534437 | CLIPA2 | CLIPB10 | 0.531411 | SPCLIP1 | CLIPB13 | 0.524655 | CLIPA7 | SP213 | 0.600605 |
| CLIPA1 | CLIPC6 | 0.448262 | CLIPA2 | CLIPB12 | 0.54302 | SPCLIP1 | CLIPB15 | 0.544791 | CLIPA8 | CLIPA9 | 0.711032 |
| CLIPA1 | CLIPC7 | 0.529729 | CLIPA2 | CLIPB13 | 0.469703 | SPCLIP1 | CLIPB4 | 0.467682 | CLIPA8 | CLIPB1 | 0.466963 |
| CLIPA1 | CLIPC9 | 0.534501 | CLIPA2 | CLIPB15 | 0.571017 | SPCLIP1 | CLIPB7 | 0.469748 | CLIPA8 | CLIPB10 | 0.624883 |
| CLIPA1 | CLIPD6 | 0.565832 | CLIPA2 | CLIPB2 | 0.508375 | SPCLIP1 | CLIPB8 | 0.463213 | CLIPA8 | CLIPB12 | 0.479497 |
| CLIPA1 | CLIPE4 | 0.521381 | CLIPA2 | CLIPB4 | 0.5483 | SPCLIP1 | CLIPC12 | 0.634349 | CLIPA8 | CLIPB13 | 0.530317 |
| CLIPA1 | CLIPE5 | 0.590239 | CLIPA2 | CLIPC12 | 0.538409 | SPCLIP1 | CLIPD1 | 0.483618 | CLIPA8 | CLIPB15 | 0.705528 |
| CLIPA1 | CLIPE10 | 0.506584 | CLIPA2 | CLIPC3 | 0.443828 | SPCLIP1 | CLIPD6 | 0.615603 | CLIPA8 | CLIPB17 | 0.518518 |
| CLIPA1 | SP213 | 0.590759 | CLIPA2 | CLIPC4 | 0.445159 | SPCLIP1 | CLIPE5 | 0.520743 | CLIPA8 | CLIPB2 | 0.709704 |
| CLIPA10 | CLIPA15 | 0.597157 | CLIPA2 | CLIPC6 | 0.490464 | SPCLIP1 | CLIPE6 | 0.471007 | CLIPA8 | CLIPB20 | 0.473831 |
| CLIPA10 | CLIPA32 | 0.457032 | CLIPA2 | CLIPC7 | 0.493746 | SPCLIP1 | SP213 | 0.682021 | CLIPA8 | CLIPB3a | 0.444043 |

| Undirected Edge | | Edge | Undirected Edge | | Edge | Undirected Edge | | Edge | Undirected Edge | | Edge Weigh |
|---|---|---|---|---|---|---|---|---|---|---|---|
| Node 1 | Node 2 | Weight | Node 1 | Node 2 | Weight | Node 1 | Node 2 | Weight | Node 1 | Node 2 | |
| CLIPA8 | CLIPB4 | 0.494932 | CLIPB10 | CLIPB13 | 0.571356 | CLIPB16 | CLIPC3 | 0.439459 | CLIPB41 | CLIPE27 | 0.55513[illegible] |
| CLIPA8 | CLIPB7 | 0.502692 | CLIPB10 | CLIPB15 | 0.518163 | CLIPB16 | SP213 | 0.491415 | CLIPB42 | CLIPC14 | 0.4930[illegible] |
| CLIPA8 | CLIPB46 | 0.460725 | CLIPB10 | CLIPB2 | 0.551191 | CLIPB17 | CLIPD1 | 0.498289 | CLIPB42 | CLIPD4 | 0.49534[illegible] |
| CLIPA8 | CLIPC12 | 0.608569 | CLIPB10 | CLIPB20 | 0.490638 | CLIPB17 | SP213 | 0.530242 | CLIPB42 | SP212 | 0.5311[illegible] |
| CLIPA8 | CLIPC3 | 0.527455 | CLIPB10 | CLIPB4 | 0.574448 | CLIPB19 | CLIPC3 | 0.545967 | CLIPB44 | CLIPB47 | 0.46990[illegible] |
| CLIPA8 | CLIPC7 | 0.512084 | CLIPB10 | CLIPB7 | 0.63209 | CLIPB19 | CLIPE10 | 0.553883 | CLIPB44 | CLIPC14 | 0.48353[illegible] |
| CLIPA8 | CLIPC9 | 0.523685 | CLIPB10 | CLIPC12 | 0.570132 | CLIPB19 | CLIPE15 | 0.5083 | CLIPB45 | CLIPE24 | 0.47829[illegible] |
| CLIPA8 | CLIPD1 | 0.486826 | CLIPB10 | CLIPC3 | 0.440854 | CLIPB19 | CLIPE17 | 0.500968 | CLIPB46 | CLIPE10 | 0.45016[illegible] |
| CLIPA8 | CLIPD6 | 0.491629 | CLIPB10 | CLIPC4 | 0.569171 | CLIPB2 | CLIPB20 | 0.467918 | CLIPB47 | CLIPE6 | 0.46469[illegible] |
| CLIPA8 | CLIPE4 | 0.625655 | CLIPB10 | CLIPC9 | 0.439449 | CLIPB2 | CLIPB4 | 0.501861 | CLIPB47 | SP217 | 0.4779[illegible] |
| CLIPA8 | CLIPE5 | 0.650957 | CLIPB10 | CLIPD6 | 0.522227 | CLIPB2 | CLIPB7 | 0.444759 | CLIPC1 | CLIPD2 | 0.48885[illegible] |
| CLIPA8 | CLIPE10 | 0.472119 | CLIPB10 | CLIPE4 | 0.512238 | CLIPB2 | CLIPB8 | 0.47336 | CLIPC1 | CLIPD4 | 0.47495[illegible] |
| CLIPA8 | SP213 | 0.561406 | CLIPB10 | CLIPE5 | 0.552177 | CLIPB2 | CLIPC12 | 0.591331 | CLIPC12 | CLIPC4 | 0.47715[illegible] |
| CLIPA9 | CLIPB1 | 0.5351 | CLIPB10 | CLIPE6 | 0.442373 | CLIPB2 | CLIPC3 | 0.572395 | CLIPC12 | CLIPC9 | 0.49488[illegible] |
| CLIPA9 | CLIPB10 | 0.549037 | CLIPB10 | CLIPE10 | 0.493045 | CLIPB2 | CLIPC9 | 0.560467 | CLIPC12 | CLIPD6 | 0.56935[illegible] |
| CLIPA9 | CLIPB12 | 0.563797 | CLIPB10 | SP213 | 0.508396 | CLIPB2 | CLIPD6 | 0.471657 | CLIPC12 | CLIPE4 | 0.57891[illegible] |
| CLIPA9 | CLIPB13 | 0.601816 | CLIPB11 | CLIPB4 | 0.50789 | CLIPB2 | CLIPE4 | 0.621129 | CLIPC12 | CLIPE5 | 0.63614[illegible] |
| CLIPA9 | CLIPB15 | 0.636404 | CLIPB11 | CLIPB5 | 0.563357 | CLIPB2 | CLIPE5 | 0.617481 | CLIPC12 | CLIPE10 | 0.51870[illegible] |
| CLIPA9 | CLIPB17 | 0.481346 | CLIPB11 | CLIPB7 | 0.454483 | CLIPB2 | CLIPE10 | 0.548883 | CLIPC12 | SP213 | 0.49343[illegible] |
| CLIPA9 | CLIPB2 | 0.631511 | CLIPB11 | CLIPB9 | 0.454442 | CLIPB2 | SP213 | 0.447453 | CLIPC13 | CLIPD20 | 0.45318[illegible] |
| CLIPA9 | CLIPB20 | 0.458981 | CLIPB11 | CLIPB47 | 0.447252 | CLIPB20 | CLIPB3a | 0.467765 | CLIPC13 | CLIPE2 | 0.47227[illegible] |
| CLIPA9 | CLIPB4 | 0.610797 | CLIPB11 | CLIPC4 | 0.46659 | CLIPB20 | CLIPB7 | 0.439429 | CLIPC13 | CLIPE22 | 0.56723[illegible] |
| CLIPA9 | CLIPB8 | 0.526467 | CLIPB12 | CLIPB2 | 0.48284 | CLIPB20 | CLIPB46 | 0.472381 | CLIPC13 | CLIPE30 | 0.52252[illegible] |
| CLIPA9 | CLIPC12 | 0.58213 | CLIPB12 | CLIPB46 | 0.473081 | CLIPB20 | CLIPC12 | 0.659822 | CLIPC13 | SP214 | 0.45093[illegible] |
| CLIPA9 | CLIPC3 | 0.65572 | CLIPB12 | CLIPC12 | 0.514818 | CLIPB20 | CLIPE10 | 0.46938 | CLIPC14 | CLIPD4 | 0.46190[illegible] |
| CLIPA9 | CLIPC4 | 0.44332 | CLIPB12 | CLIPC6 | 0.492452 | CLIPB20 | SP213 | 0.461177 | CLIPC14 | SP217 | 0.48883[illegible] |
| CLIPA9 | CLIPC9 | 0.614199 | CLIPB12 | CLIPE4 | 0.541108 | CLIPB3a | CLIPC12 | 0.449619 | CLIPC3 | CLIPC4 | 0.48120[illegible] |
| CLIPA9 | CLIPD6 | 0.502667 | CLIPB12 | CLIPE5 | 0.514928 | CLIPB3a | CLIPC9 | 0.493135 | CLIPC3 | CLIPC9 | 0.57356[illegible] |
| CLIPA9 | CLIPE4 | 0.679912 | CLIPB12 | CLIPE10 | 0.566954 | CLIPB3a | CLIPE2 | 0.475653 | CLIPC3 | CLIPD6 | 0.48217[illegible] |
| CLIPA9 | CLIPE5 | 0.706191 | CLIPB13 | CLIPB15 | 0.578047 | CLIPB3b | CLIPC9 | 0.452635 | CLIPC3 | CLIPD11 | 0.52484[illegible] |
| CLIPA9 | CLIPE6 | 0.510638 | CLIPB13 | CLIPB2 | 0.507723 | CLIPB36 | CLIPB4 | 0.487818 | CLIPC3 | CLIPE4 | 0.54644[illegible] |
| CLIPA9 | CLIPE10 | 0.626903 | CLIPB13 | CLIPB20 | 0.480796 | CLIPB36 | CLIPC4 | 0.496377 | CLIPC3 | CLIPE5 | 0.56694[illegible] |
| CLIPA9 | SP213 | 0.710012 | CLIPB13 | CLIPB4 | 0.573626 | CLIPB36 | CLIPC6 | 0.498589 | CLIPC3 | CLIPE10 | 0.47481[illegible] |
| CLIPA31 | CLIPA32 | 0.525836 | CLIPB13 | CLIPB8 | 0.495014 | CLIPB36 | CLIPD6 | 0.486325 | CLIPC3 | SP213 | 0.6185[illegible] |
| CLIPA31 | CLIPB45 | 0.451378 | CLIPB13 | CLIPC12 | 0.480333 | CLIPB36 | CLIPE1 | 0.482061 | CLIPC4 | CLIPC6 | 0.52104[illegible] |
| CLIPA31 | CLIPD8 | 0.455049 | CLIPB13 | CLIPC3 | 0.498223 | CLIPB36 | CLIPE18 | 0.543831 | CLIPC4 | CLIPD6 | 0.47018[illegible] |
| CLIPA31 | CLIPD12 | 0.476681 | CLIPB13 | CLIPC7 | 0.495973 | CLIPB4 | CLIPB7 | 0.470033 | CLIPC4 | CLIPE5 | 0.49927[illegible] |
| CLIPA31 | CLIPD13 | 0.460632 | CLIPB13 | CLIPC9 | 0.51668 | CLIPB4 | CLIPB8 | 0.551641 | CLIPC6 | CLIPC7 | 0.51349[illegible] |
| CLIPA31 | CLIPE18 | 0.56431 | CLIPB13 | CLIPD6 | 0.545286 | CLIPB4 | CLIPC3 | 0.653557 | CLIPC6 | CLIPD6 | 0.53709[illegible] |
| CLIPA31 | CLIPE19 | 0.53774 | CLIPB13 | CLIPE5 | 0.513268 | CLIPB4 | CLIPC4 | 0.660585 | CLIPC6 | CLIPE23 | 0.44474[illegible] |
| CLIPA31 | CLIPE21 | 0.681154 | CLIPB13 | CLIPE6 | 0.473291 | CLIPB4 | CLIPC6 | 0.53498 | CLIPC7 | CLIPD6 | 0.45038[illegible] |
| CLIPA31 | CLIPE25 | 0.549225 | CLIPB13 | SP213 | 0.545259 | CLIPB4 | CLIPC9 | 0.573478 | CLIPC9 | CLIPD6 | 0.51604[illegible] |
| CLIPA32 | CLIPD7 | 0.479325 | CLIPB14 | CLIPB20 | 0.481731 | CLIPB4 | CLIPD6 | 0.553273 | CLIPC9 | CLIPE4 | 0.48778[illegible] |
| CLIPA32 | CLIPD13 | 0.499356 | CLIPB15 | CLIPB17 | 0.52766 | CLIPB4 | CLIPE5 | 0.520863 | CLIPC9 | CLIPE5 | 0.49452[illegible] |
| CLIPA32 | CLIPD20 | 0.633826 | CLIPB15 | CLIPB2 | 0.591254 | CLIPB4 | SP213 | 0.549827 | CLIPC9 | CLIPE10 | 0.52276[illegible] |
| CLIPA32 | CLIPE2 | 0.473374 | CLIPB15 | CLIPB20 | 0.473557 | CLIPB5 | CLIPC10 | 0.458211 | CLIPC9 | CLIPE17 | 0.53275[illegible] |
| CLIPA32 | CLIPE18 | 0.452995 | CLIPB15 | CLIPB3a | 0.499664 | CLIPB7 | CLIPD6 | 0.513117 | CLIPC9 | SP213 | 0.4867[illegible] |
| CLIPA32 | CLIPE21 | 0.460034 | CLIPB15 | CLIPB4 | 0.547173 | CLIPB7 | CLIPE6 | 0.46682 | CLIPD1 | SP213 | 0.47980[illegible] |
| CLIPB1 | CLIPB10 | 0.443064 | CLIPB15 | CLIPB8 | 0.524166 | CLIPB7 | SP213 | 0.504414 | CLIPD2 | CLIPD7 | 0.5390[illegible] |
| CLIPB1 | CLIPB13 | 0.587992 | CLIPB15 | CLIPC12 | 0.612652 | CLIPB8 | CLIPC10 | 0.475007 | CLIPD2 | CLIPD12 | 0.44655[illegible] |
| CLIPB1 | CLIPB15 | 0.495944 | CLIPB15 | CLIPC3 | 0.538231 | CLIPB8 | CLIPC3 | 0.46729 | CLIPD2 | CLIPD22 | 0.52668[illegible] |
| CLIPB1 | CLIPB2 | 0.473062 | CLIPB15 | CLIPC7 | 0.477959 | CLIPB8 | CLIPC7 | 0.528226 | CLIPD3 | CLIPD8 | 0.58951[illegible] |
| CLIPB1 | CLIPB4 | 0.7508 | CLIPB15 | CLIPC9 | 0.587713 | CLIPB8 | CLIPC9 | 0.488918 | CLIPD3 | CLIPD12 | 0.57955[illegible] |
| CLIPB1 | CLIPB8 | 0.500172 | CLIPB15 | CLIPD1 | 0.528807 | CLIPB41 | CLIPD12 | 0.476506 | CLIPD3 | CLIPD13 | 0.45591[illegible] |
| CLIPB1 | CLIPC3 | 0.567804 | CLIPB15 | CLIPD6 | 0.613819 | CLIPB41 | CLIPD13 | 0.542319 | CLIPD3 | CLIPD14 | 0.58453[illegible] |
| CLIPB1 | CLIPC4 | 0.56451 | CLIPB15 | CLIPE4 | 0.516527 | CLIPB41 | CLIPD22 | 0.472588 | CLIPD3 | CLIPD22 | 0.49890[illegible] |
| CLIPB1 | CLIPC6 | 0.563724 | CLIPB15 | CLIPE5 | 0.517338 | CLIPB41 | CLIPE2 | 0.52552 | CLIPD3 | CLIPE25 | 0.56447[illegible] |
| CLIPB1 | CLIPC9 | 0.469454 | CLIPB15 | CLIPE10 | 0.504266 | CLIPB41 | CLIPE23 | 0.485416 | CLIPD4 | CLIPE13 | 0.44848[illegible] |
| CLIPB1 | CLIPD6 | 0.509436 | CLIPB15 | SP213 | 0.537243 | CLIPB41 | CLIPE25 | 0.495337 | CLIPD4 | CLIPE23 | 0.53021[illegible] |

41

| Undirected Edge | | Edge |
| --- | --- | --- |
| **Node 1** | **Node 2** | **Weight** |
| CLIPD6 | CLIPE4 | 0.459775 |
| CLIPD6 | SP213 | 0.463827 |
| CLIPD7 | CLIPD12 | 0.654158 |
| CLIPD7 | CLIPD13 | 0.567557 |
| CLIPD7 | CLIPD22 | 0.464095 |
| CLIPD8 | CLIPD12 | 0.552431 |
| CLIPD8 | CLIPD13 | 0.509795 |
| CLIPD8 | CLIPD14 | 0.510447 |
| CLIPD8 | CLIPE25 | 0.561585 |
| CLIPD12 | CLIPD13 | 0.642623 |
| CLIPD12 | CLIPD22 | 0.68357 |
| CLIPD12 | CLIPE2 | 0.51882 |
| CLIPD12 | CLIPE21 | 0.452946 |
| CLIPD12 | CLIPE25 | 0.510575 |
| CLIPD13 | CLIPD22 | 0.600007 |
| CLIPD13 | CLIPE25 | 0.537471 |
| CLIPD20 | CLIPD22 | 0.457193 |
| CLIPD20 | CLIPE2 | 0.546176 |
| CLIPD20 | CLIPE23 | 0.450824 |
| CLIPD20 | SP214 | 0.495363 |
| CLIPD22 | CLIPE13 | 0.48648 |
| CLIPD22 | CLIPE23 | 0.447239 |
| CLIPD22 | CLIPE25 | 0.512554 |
| CLIPE2 | CLIPE19 | 0.479651 |
| CLIPE2 | CLIPE23 | 0.459752 |
| CLIPE2 | CLIPE30 | 0.489239 |
| CLIPE2 | SP214 | 0.526589 |
| CLIPE4 | CLIPE5 | 0.664287 |
| CLIPE4 | CLIPE10 | 0.621539 |
| CLIPE4 | SP213 | 0.447493 |
| CLIPE5 | CLIPE10 | 0.697331 |
| CLIPE5 | SP213 | 0.448605 |
| CLIPE6 | SP213 | 0.656303 |
| CLIPE9 | CLIPE34 | 0.522446 |
| CLIPE10 | CLIPE17 | 0.564346 |
| CLIPE11 | CLIPE29 | 0.503081 |
| CLIPE12 | CLIPE16 | 0.533954 |
| CLIPE12 | CLIPE17 | 0.483288 |
| CLIPE13 | CLIPE23 | 0.487339 |
| CLIPE13 | SP212 | 0.46094 |
| CLIPE16 | CLIPE17 | 0.580812 |
| CLIPE17 | CLIPE29 | 0.515297 |
| CLIPE18 | CLIPE30 | 0.487132 |
| CLIPE19 | CLIPE25 | 0.490853 |
| CLIPE21 | CLIPE23 | 0.533919 |
| CLIPE22 | CLIPE28 | 0.505625 |
| CLIPE22 | CLIPE34 | 0.502861 |
| CLIPE23 | CLIPE25 | 0.579379 |
| CLIPE24 | CLIPE27 | 0.497751 |

**Online suppl. Table S4. ESI-MS analysis of the CLIPB4-SRPN2 complex**

| Protein | AGAP # (AgamP4.13) | MS matched peptide counts | Matched peptide sequence | Mascot score | Total score |
|---|---|---|---|---|---|
| rSRPN2 | AGAP006911 | 15 | R.QNEFDLMFVK.E | 18.9 | 1027.8 |
| | | | K.ILLTLIYEASDTSFGNAVSNTK.R | 42.6 | |
| | | | R.ELSSVIQNDNIDHTR.S | 30.1 | |
| | | | K.VSYSNPTQTAATINNWVSEHTNGR.L | 103.6 | |
| | | | R.EIVTPDSLEGAVITLVNVIYFK.G | 113.9 | |
| | | | R.GKPTNAQYMEQNGQFYYDNSADLGAQILR.L | 109.4 | |
| | | | K.LAMYFILPNPDNTVNQVLDR.I | 69.6 | |
| | | | K.LAMYFILPNPDNTVNQVLDR.I | 54.1 | |
| | | | K.FKFDFSEQLNEPLQQVGIR.E | 89.9 | |
| | | | K.FDFSEQLNEPLQQVGIR.E | 79.5 | |
| | | | R.EIFSQNASLPLLAR.G | 67.7 | |
| | | | K.AGITINELGSEAYAATEIQLVNK.F | 43.8 | |
| | | | K.AGITINELGSEAYAATEIQLVNK.F | 63.5 | |
| | | | K.FGGDGVQIFNANRPFIFFIEDETLGTMLFAGK.I | 59.6 | |
| | | | K.FGGDGVQIFNANRPFIFFIEDETLGTMLFAGK.I | 81.6 | |
| proCLIPB4$_{Xa}$ | AGAP003250 | 2 | K.IDEFPWTALIEYEKPNGR.F | 76.2 | 121.2 |
| | | | R.LGEWDLSSTTDQEDDFYADAPIDLDIEK.I | 45.0 | |

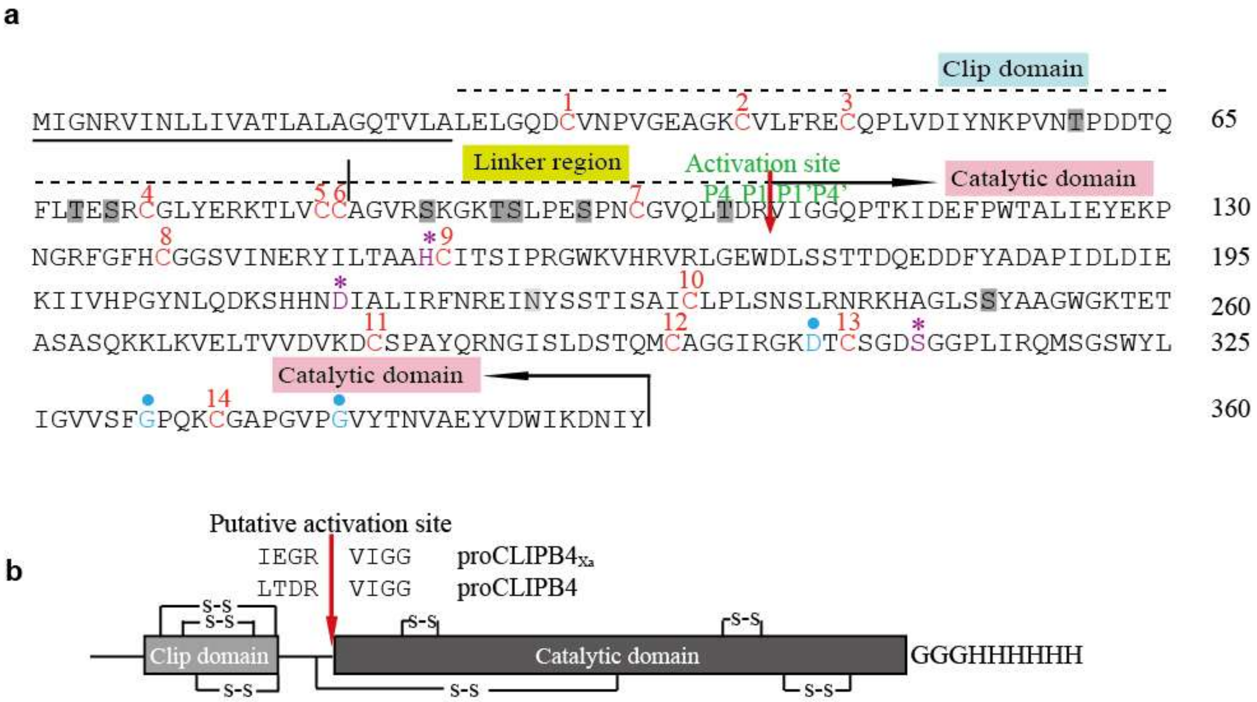


**Online suppl. Fig. S1. Sequence feature analysis of CLIPB4.** (**a**) Amino acid residues of CLIPB4. CLIPB4 consists of one clip domain and one catalytic domain (denoted above the amino acid sequence). The predicted signal peptide is underlined. Conserved cysteine residues are shown in red and indicated with red numbers on the basis of determined structure of *M. sexta* PAP2. The catalytic residues (His, Asp, Ser) of CLIPB4 are shown in purple and purple asterisks. The amino acid residues that determine the primary specificity of CLIPB4 are marked with blue circles. The predicted activation site is highlighted with red arrow. There are potentially 1 *N*-linker and 9 *O*-linker glycosylation sites in CLIPB4, which are lightly and heavily shaded, respectively. (**b**) Schematic representation of recombinant CLIPB4. The conserved Cys residues in (**a**) are connected by disulfide bonds (marked with "s-s"). In the mutant proCLIPB4$_{Xa}$, its activation residues were manually replaced with IEGR to permit the activation by bovine Factor Xa (denoted with red arrow).

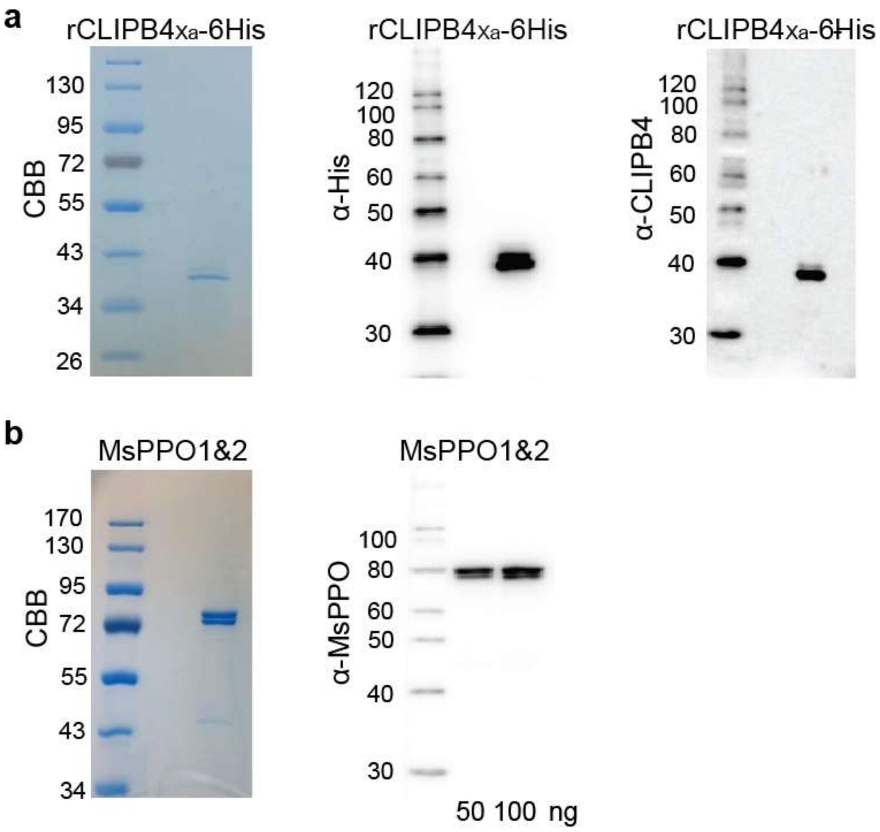


**Online suppl. Fig. S2. Recombinant and purified proteins in this study.** Detection of purified recombinant proCLIPB4$_{Xa}$-6His and *Ms*PPO1&2 (from *M. sexta* $5^{th}$ instar larval hemolymph) by reducing SDS-PAGE, Coomassie brilliant blue staining (CBB) and immunoblot. (**a**) Recombinant proCLIPB4$_{Xa}$-6His. 300 ng for CBB, 150 ng for mouse anti-His antibody, and 20 ng for rabbit anti-CLIPB4 antibody. (**b**) Purified *Ms*PPO1&2 from *M. sexta* larvae. 1 μg for CBB; 50 and 100 ng for rabbit anti-*Ms*PPO1&2 antibody.

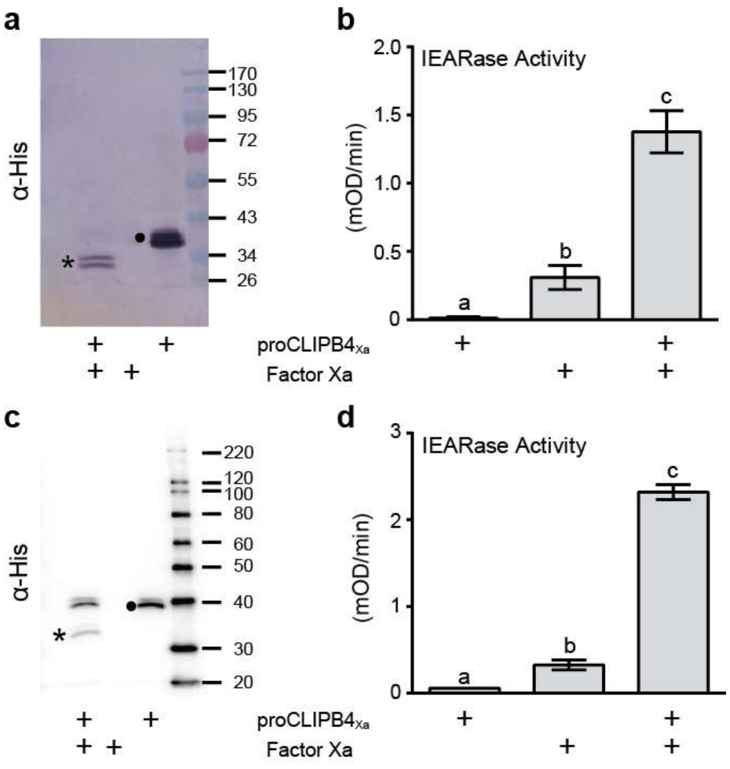


**Fig. S3. Activation of recombinant proCLIPB4$_{Xa}$ by Factor Xa.**

To activate recombinant proCLIPB4$_{Xa}$, 2.5 µg zymogen was mixed with 1 µg bovine Factor-Xa (1 µg/µl, New England Biolabs), supplemented to 25 µl with Factor-Xa activation buffer (20 mM Tris, 100 mM NaCl, 2 mM $CaCl_2$, pH 8.0). Two negative controls were set up in parallel by replacing proCLIPB4$_{Xa}$ and Factor-Xa with equal volume of activation buffer, respectively. Reactions were incubated either at 37°C (**a,b**) or RT (**c,d**) overnight. To confirm activation cleavage, we subjected each activation mixture to SDS-PAGE and immunoblot using mouse anti-His antibody, loading volumes equivalent to 154 ng (**a**) or 150 ng (**c**) of proCLIPB4$_{Xa}$. *Circle*, recombinant proCLIPB4$_{Xa}$ zymogen; *asterisk*, catalytic domain of proCLIPB4$_{Xa}$. To confirm protease activity of Factor Xa-activated CLIPB4$_{Xa}$, a small volume of each mixture, equivalent to 91 ng (**b**) or 100 ng (**d**) of proCLIPB4$_{Xa}$ was added in a 96-well plate. 200 µl 50 µM substrate IEAR*p*NA was added per well to examine amidase activity (N-acetyl-benzoyl-Ile-Glu-Ala-Arg-p-nitroanilide [bioWORLD] in 0.1 M Tris, 0.1 M NaCl, 5 mM $CaCl_2$, pH 8.0). Absorbance change was monitored at 405 nm in EPOCH 2 microplate reader (BioTek), and one unit of amidase activity was defined as ΔA405/min = 0.001. Three technical replicates were performed each time. The bars represent mean±1 SD (n=3). Statistically significant differences are marked with different letters (one-way ANOVA followed by Newman-Keuls test, $P<0.05$).

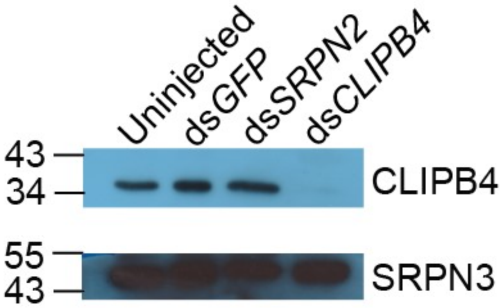


**Online suppl. Fig. S4. ds*CLIPB4* depletes CLIPB4 in *An. gambiae* hemolymph.** Hemolymph was collected by proboscis clipping from 8 female mosquitoes 4 days post injection. Samples were subjected to SDS-PAGE and immunoblot detection with rabbit anti-CLIPB4 and rabbit anti-SRPN3 antibodies. SRPN3 antibody was stripped before CLIPB4 antibody probing.

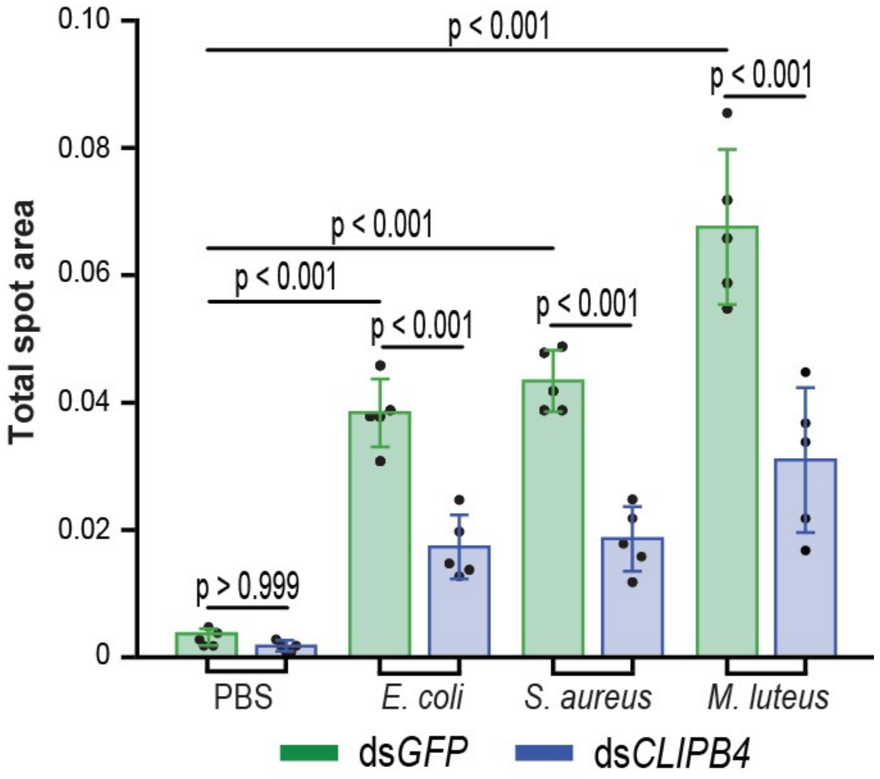


**Online suppl. Fig. S5. *CLIPB4* is required for melanization after microbial challenge.** 60 h post dsRNA treatment, each mosquito was challenged by systemic injection (50.6 nl) of PBS, *E. coli* ($OD_{600}$=1.5), *S. aureus* ($OD_{600}$=1.5) or *M. luteus* ($OD_{600}$=0.4). Anal droplets were collected on a clean filter paper within 12 h post bacterial challenge. Five biological replicates were performed with 50 females per treatment group in each replicate. The bars represent mean ± 1 SD, and P values were calculated with one-way ANOVA.

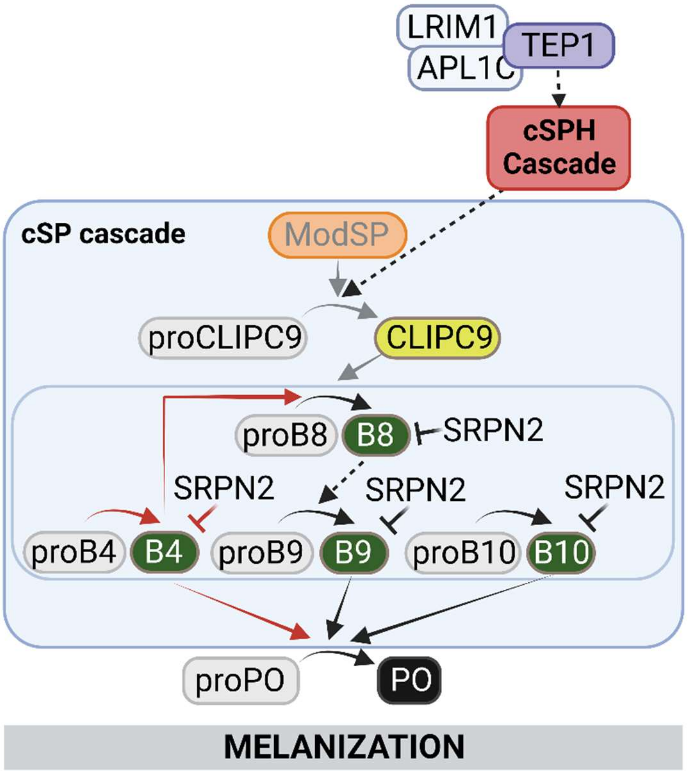


**Online suppl. Fig. S6. Model of proPO activation in *An. gambiae***. Melanization in *An. gambiae* is regulated by at least four cSPs in the CLIPB family (green) and one cSP in the CLIPC family (yellow). These CLIPs circulate in the hemolymph of adult female mosquitoes as zymogens and are cleaved in response to infection. We hypothesize that a modular serine protease (orange) cleaves and activates CLIPC9, which in turn cleaves one or more of the four downstream proCLIPBs. In addition, the cSP cascade is regulated by the action of TEP1 (purple) and a cSPH cascade (red), consisting of SPCLIP1, CLIPA28, and CLIPA8. Solid arrows indicate direct biochemical interactions and proteolytic activation, and black lines indicating direct inhibition by serpins. Dashed arrows represent interactions based on genetic evidence. Red lines, this study; black lines, published previously; gray lines, data available in other insect species only. Figure was created with BioRender.com.
